# cylcop: An R Package for Circular-Linear Copulae with Angular Symmetry

**DOI:** 10.1101/2021.07.14.452253

**Authors:** Florian H. Hodel, John R. Fieberg

## Abstract

The R package **cylcop** extends the **copula** package to allow modeling of correlated circular-linear random variables using copulae that are symmetric in the circular dimension. We present and derive several new circular-linear copulae with this property and demonstrate how they can be implemented in the **cylcop** package to model animal movements in discrete time. The package contains methods for estimating copulae parameters, plotting probability density and cumulative distribution functions, and simulating data.

## 1. Introduction

Circular-linear data often arise when modeling outcomes with a directional component (e.g., pollutant concentrations with accompanying wind directions; García-Portugués, Crujeiras, and González-Manteiga 2013). They are also common in the context of modeling animal movement as a time series of discrete steps characterized by step lengths (distance between locations) and turn angles (change in direction from the previous step) (Hodel and Fieberg 2021). Step lengths and turn angles are often correlated, with animals cycling between directed movements with long steps and turn angles near 0 when traveling and short steps with large turn angles when foraging. Movement data are also somewhat unique, in that the circular variable (turn angle) is frequently symmetric, meaning that an animal has no inherent preference to turn left or right. This correlation and symmetry can be captured using a copula, which is a probability distribution with uniform marginals on the interval [0, 1]. In this paper, we will only focus on bivariate copulae, but our work can in principle be extended to higher dimensions. We derive several copulae that can be used to model symmetric circular-linear data and and present their implementation in a new R package **cylcop** (R Core Team 2021), freely available from the Comprehensive R Archive Network (CRAN) at https://CRAN.R-project.org/package=cylcop. The **cylcop** package extends the classes and methods available in the **copula** package (Hofert, Kojadinovic, Maechler, and Yan 2020; Jun Yan 2007; Ivan Kojadinovic and Jun Yan 2010; Marius Hofert and Martin Mächler 2011; Hofert, Kojadinovic, Maechler, and Yan 2018) and also builds on the **circular** package (Agostinelli and Lund 2017). We describe the derivation and implementation of new circular-linear copulae here and provide an introduction to circular-linear copulae and their application to movement data elsewhere (Hodel and Fieberg 2021). Finally, we also provide an app that allows for interactive visualization and exploration of all copulae introduced in this paper which can be accessed at https://cylcop.shinyapps.io/cylcop-graphs/.

## 2. Introduction to copulae

A bivariate copula *C*(*u, v*) is a cumulative distribution function (CDF) with uniform marginals and its domain restricted to the unit square, (*u, v*) ∈ [0, 1]^2^. Using general properties of CDFs, we can therefore mathematically define a copula to be any function *C*(*u, v*) : [0, 1]^2^ → [0, 1] that fulfills

1. *C*(*u*, 0) = *C*(0, *v*) = 0, ∀*u, v* ∈ [0, 1].
2. *C*(*u*, 1) = *u* and *C*(1, *v*) = *v*, ∀*u*, *v* ∈ [0, 1].
3. *C*(*u, v*) is 2-increasing.

2-increasing means that for any rectangle [*u, u*′] × [*v, v*′] with *u, u*′, *v, v*′ ∈ [0, 1], (*u* ≤ *u*′) and (*v* ≤ *v*′), the C-volume *V_C_* (i.e., the probability of (*U, V*) taking a value from the subset of the unit square described by the rectangle) is non-negative

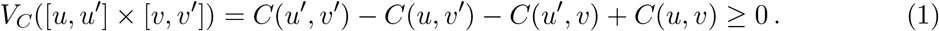

In combination with the first property, i.e., a copula being a grounded function, this means that a copula is non-decreasing in both arguments.

For any two random variables, *X* and *Y*, with strictly increasing and continuous CDFs, *F_X_*(*x*) = *P*(*X* ≤ *x*) and *F_Y_*(*y*) = *P*(*Y* ≤ *y*), and their inverses, 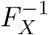 and 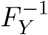 (defined as 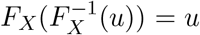, and 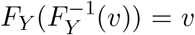, ∀*u, v* ∈ [0, 1]), we can find a joint CDF where the correlation structure is captured by the chosen copula. Let *U* and *V* be uniformly distributed random variables. Then, using the probability integral transform, we have

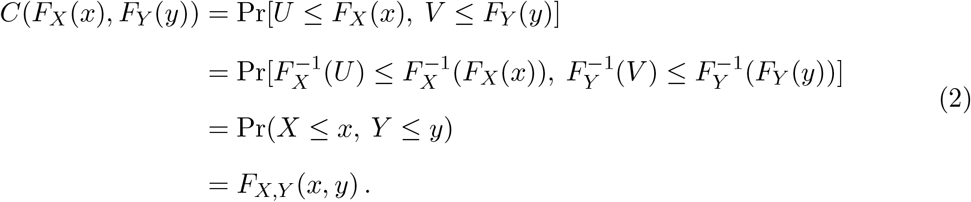

This is called Sklar’s theorem (Sklar 1959).

The probability density function (PDF), *f_X,Y_*(*x, y*), of that joint distribution is

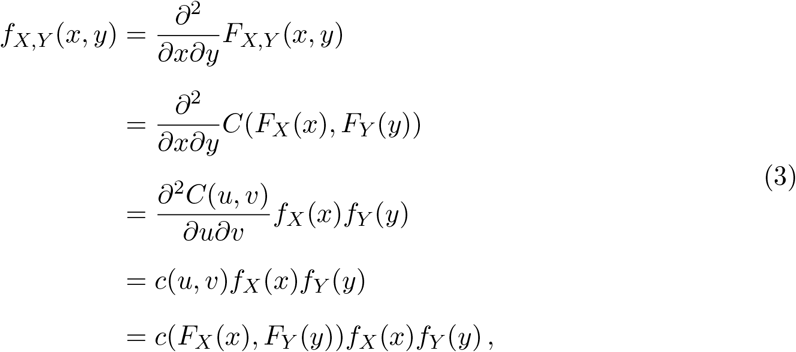

where we made the substitutions *F_X_*(*x*) = *u* and *F_Y_*(*y*) = *v*, and *f_X_*(*x*) and *f_Y_*(*y*) denote the marginal PDFs of *X* and *Y*, respectively. *c*(*u, v*) is the copula density, i.e., a PDF of a distribution with uniform marginals corresponding to the CDF *C*(*u, v*). Hereafter, we will use “copula” to refer to *C*(*u, v*) and “copula density” to refer to its derivative, 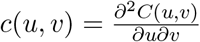. Besides calculating PDFs and CDFs, we also need to be able to draw samples from joint distributions. This can be done by drawing first a sample from a copula, which can then be transformed via the probability integral transform using the inverses of the marginal CDFs to a sample from the joint distribution. A common approach to sample copulae uses their conditional distributions (Johnson 2013)

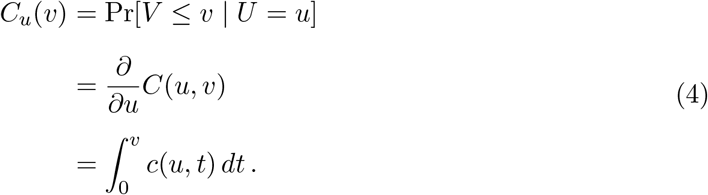

We can now take 2 independent samples from a uniform distribution, *u* and *z*, make the transformation 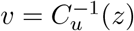, and thus obtain a sample (*u, v*) from the copula *C*(*u, v*). Finally, we can generate a sample from a joint distribution of a pair of random variables with the correlation structure described by that copula via 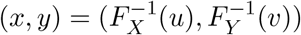.

Since a copula is a grounded, 2-increasing function, we know that firstly, *C*(*u, v*) ≤ *C*(*u*, 1) = *u* and *C*(*u, v*) ≤ *C*(1, *v*) = *v* and secondly, *V_C_*([*u*, 1] × [*v*, 1]) = 1 – *u* – *v* + *C*(*u, v*) ≥ 0 and *C*(*u, v*) ≥ 0. From the first statement, we can derive the upper (*M*(*u, v*)) and from the second the lower (*W*(*u, v*)) bound of any copula

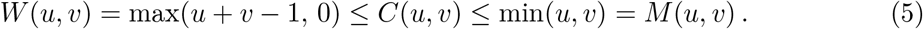

*W* and *M* are called the Fréchet–Hoeffding bounds (Fréchet 1935; Hoeffding 1940; Fréchet 1951) and are themselves copulae (at least in the bivariate case). Not all copulae can describe any degree of correlation and attain both bounds. Finally, the product copula Π(*u, v*) = *uv*, corresponds to independent random variables. For every copula introduced in this paper, we will derive the CDF, PDF, conditional copula, and Fréchet–Hoeffding bounds.

### 2.1. Empirical copula

With a sample of *n* independent and identically distributed (i.i.d.) bivariate observations (*x_i_, y_i_*) drawn from a joint distribution, we can calculate the empirical marginal CDFs

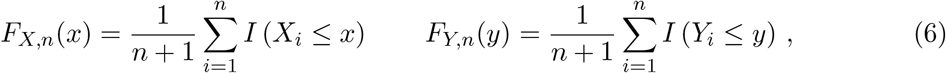

with *I* the indicator function. Transforming the observations (*x_i_, y_i_*) with the empirical marginal CDFs gives us the so-called pseudo-observations (*F_X,n_*(*x_i_*), *F_Y,n_*(*y_i_*)). Assuming continuous data, and hence a negligible probability for ties, the pseudo-observations are just the ranks (*r_i_* and *s_i_*) of the data divided by a normalization constant, 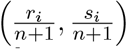. Note that we divide by *n* + 1 and not *n* to have all pseudo-observations inside the open unit square. Doing so helps avoid boundary effects and issues arising when calculating the density. The pseudo-observations can be seen as values drawn from some empirical copula, the CDF of which we can calculate as (Rüschendorf 1976; Deheuvels 1979; Genest and Favre 2007):

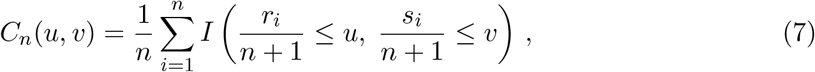

The empirical copula density is also easily accessible, but of less importance for the present work. When one of the random variables is circular, we can follow the same procedure, but we will need to define a reference point to determine the ranks of the circular variable. Throughout this paper, we will choose *–π* as our reference.

### 2.2. Copula symmetry and rotations

The survival function 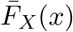 of a random variable *X* is defined as

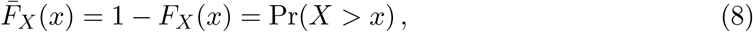

and the joint survival function of two random variables *X* and *Y* can be written using de Morgan’s laws as

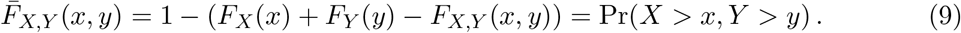

Using 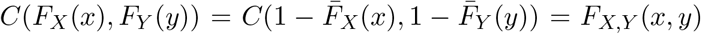, we can rewrite the above equation as

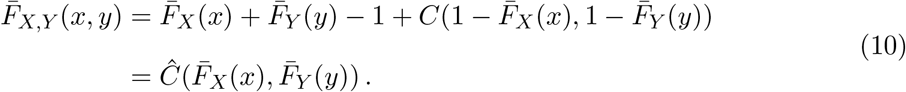

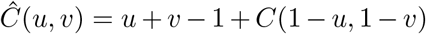 is called the survival copula and is itself a copula Nelsen (2006).

If we reflect a copula density *c*(*u, v*) with respect to the lines *u* = 0.5 and *v* = 0.5, we obtain the function *c_π_* = *c*(1 – *u*, 1 – *v*). This is commonly called “rotation” (by 180 degrees) in the copula community, e.g., in Hofert *et al*. (2018), (hence the subscript *π*). The corresponding distribution function can be found by integrating

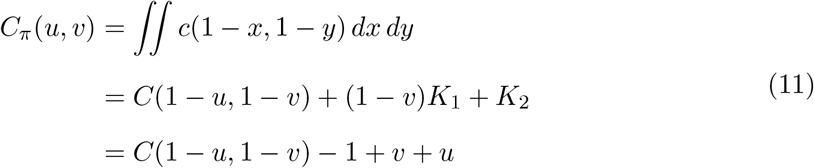

where we determined the integration constants *K*_1_ = –1 and *K*_2_ = *u* from the boundary conditions *v* = *C_π_*(1, *v*) and *u* = *C_π_*(*u*, 1). Note that this is the equation of the survival copula of *C* and thus, 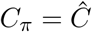. I.e., we have shown that the survival copula can be obtained by rotating the copula density by 180 degrees. “Rotations” by 90 and 270 degrees also lead to valid copulae, which are denoted by *C*_0.5π_ = *v* – *C*(1 – *u, v*) and *C*_1.5π_ = *u* – *C*(*u*, 1 – *v*). They correspond to orthogonal reflections of the copula density along the line *u* = 0.5 and *v* = 0.5, respectively.

A (continuous) random variable *X* is symmetric about *a* if its distribution fulfills

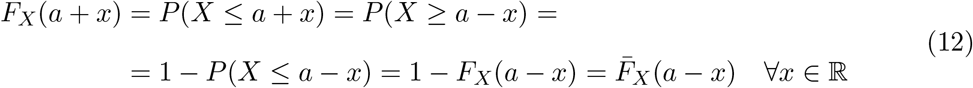

which means that *F_X_*(*a* + *x*) + *F_X_*(*a* – *x*) = 1. Hence, it must hold for the density function (if *X* admits a density) that *f_X_*(*a* + *x*) = *f_X_*(*a* – *x*). A pair of random variables (*X, Y*) is marginally symmetric about a point (*a, b*) if *X* is symmetric about *a* and *Y* is symmetric about *b*. (*X, Y*) is furthermore radially symmetric about (*a, b*) iff

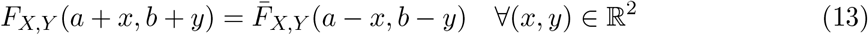

which again means that it must hold for the density function that *f_X,Y_*(*a* + *x,b* + *y*) = *f_X,Y_*(*a* – *x,b* – *y*). Radial symmetry implies marginal symmetry. From the statements about survival copulae and reflected copula densities made above, we can derive that a pair of random variables that is marginally symmetric is radially symmetric iff its copula density is radially symmetric about (0.5, 0.5), i.e.,

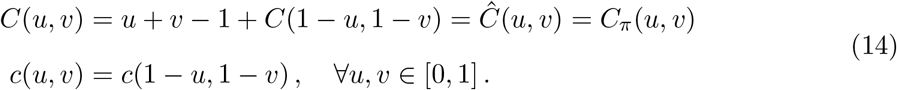

In the following sections, we will often require that pairs of random variables be symmetric in *X* for any value of *Y*. With a symmetric marginal distribution of *X*, this means that the copula must be symmetric in *u* about *u* = 0.5 and

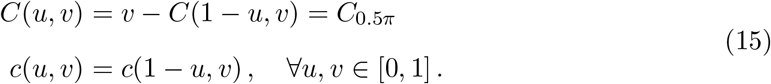

## 3. Circular distributions and their implementation

Let Θ be a continuous circular random variable with support on the unit circle, 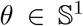. There are two ways to “unwrap” the density and distribution function. This seems like a minor point, but, in the authors’ experience, it can lead to confusion. We will elucidate the issue using the von Mises distribution, one of the most commonly used distributions, as an example. Functions giving e.g., the von Mises CDF behave differently in different program packages, and since the **cylcop** package makes heavy use of the **circular** package (Agostinelli and Lund 2017), we will take a closer look at that particular implementation.

The von Mises PDF is usually defined as a function

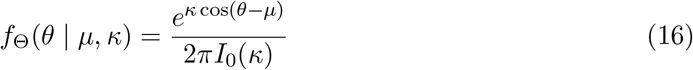

with parameters 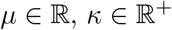 and support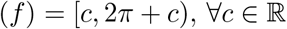. This PDF fulfills the boundary condition

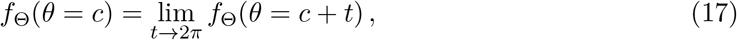

and the corresponding CDF is

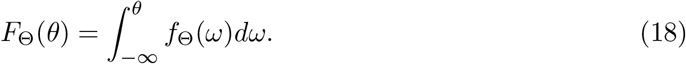

The von Mises PDF and CDF are implemented this way, e.g., in Wolfram Mathematica (Wolfram Research, Inc. 2020) (see Figure 1).

**Figure 1:**
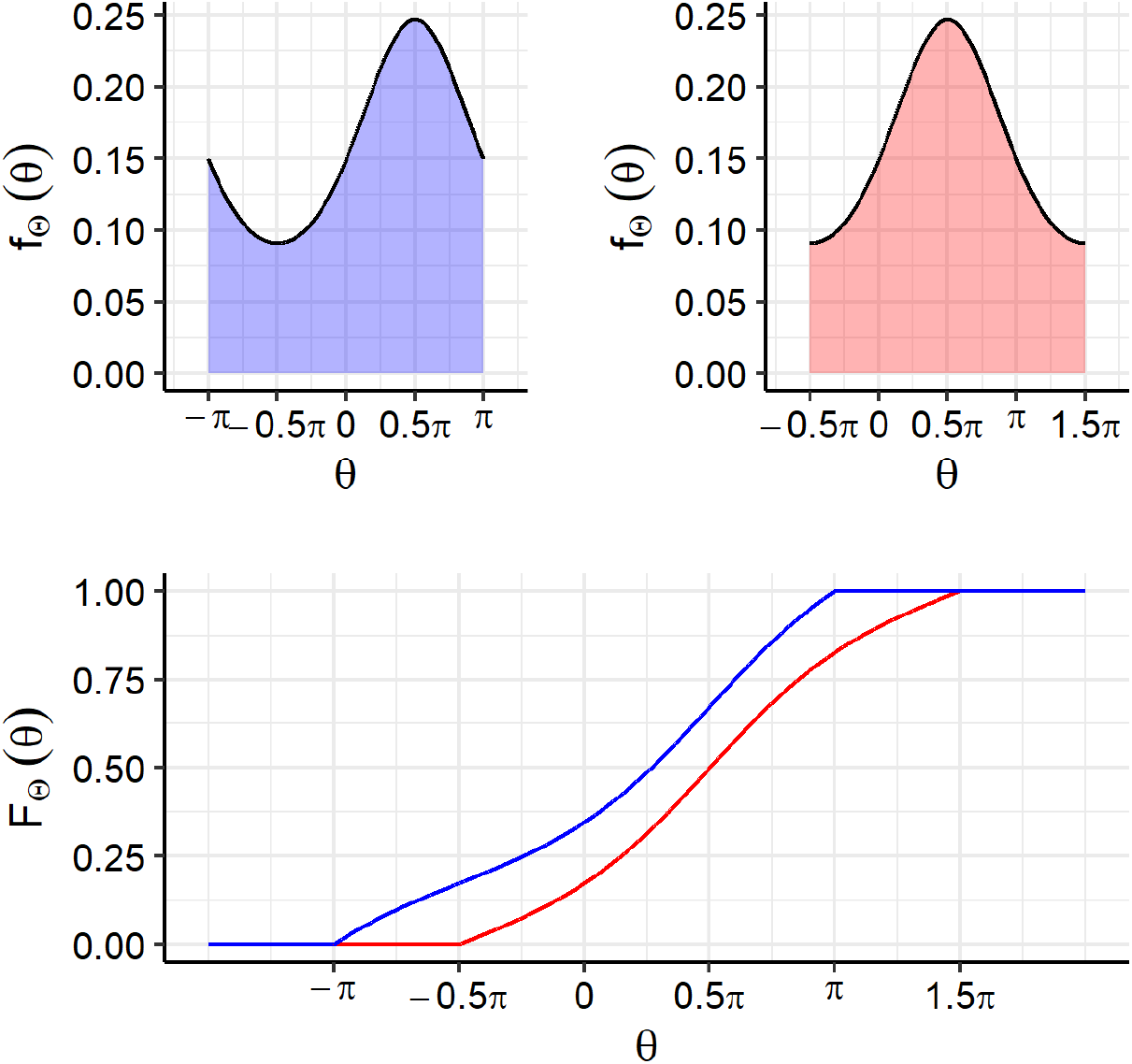
Top: PDF of a von Mises distribution with *μ* = 0.5*π* and *κ* = 0.5. Left: *c* = –*π*, right: *c* = –0.5*π*. Bottom: The corresponding CDFs.

Alternatively, we can allow the angle to assume any real value, 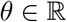, and have a PDF with support on the entire real line (which is done e.g., in Mardia and Jupp (2000)). To distinguish it from the “restricted” von Mises density above, we will call it *g*_Θ_(*θ* | *μ, κ*), the only difference to *f*_Θ_(*θ* | *μ, κ*) being that 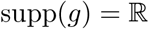, which leads to a periodic PDF with

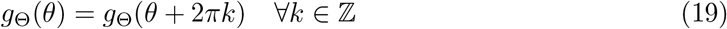

and *g*_Θ_(*θ*) integrating to one over any interval of length 2*π*. To actually calculate the periodic CDF,

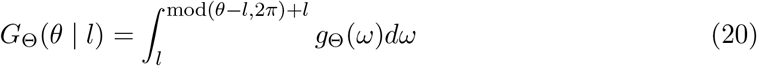

we must choose a lower limit of integration *l*. *g*(*θ*) and *G*(*θ* | *l*) correspond to the functions `pvonmises()` and `dvonmises()`, in the **circular** package (Agostinelli and Lund 2017) (see Figure 2).

**Figure 2:**
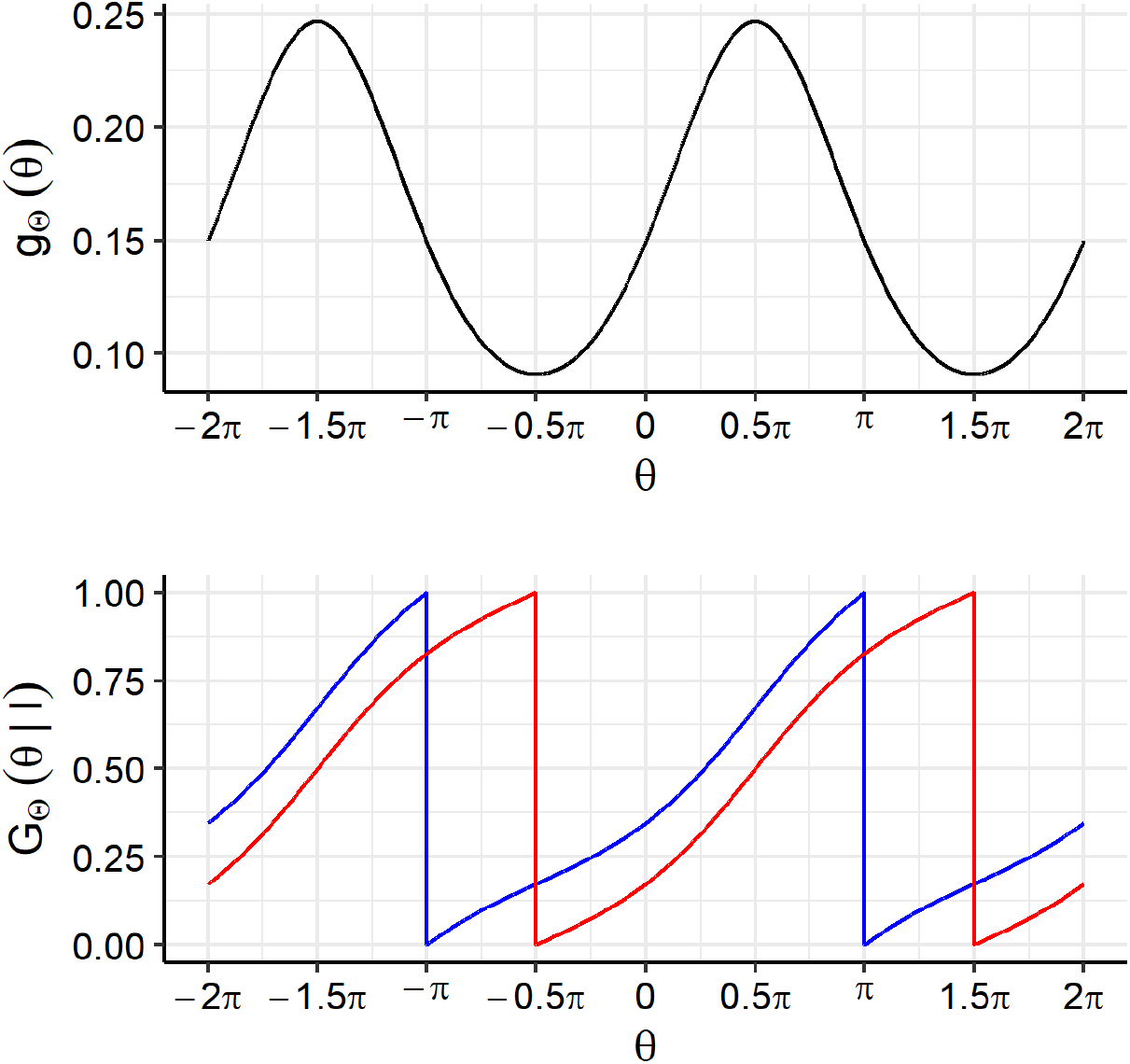
Top: PDF of a von Mises distribution with *μ* = 0.5*π* and *κ* = 0.5. Bottom: The corresponding CDFs. Blue: *l* = –*π*, red: *l* = –0.5*π*.

In summary, we can either allow a circular random variable to have the entire real line as support, which leads to periodic density and distribution functions (which is done e.g., in Mardia and Jupp (2000) and is implemented in the **circular** package), or we restrict the support to an interval of 2*π*, and have densities that need to fulfill the boundary condition in Equation 17. The two approaches of “unwrapping” a circular random variable are of course equivalent. In the context of copulae and turning angles, it is however more natural to think of *θ* as being restricted to the interval [–*π,π*).

### 3.1. Fitting circular distributions

We have implemented a function to fit parameters of certain circular distributions to a set of angles in [–*π,π*) using a maximum likelihood approach.

~~~
*R> fit_angle(theta,
+    parametric = c(“vonmises”, “wrappedcauchy”, “mixedvonmises”, FALSE),
+    bandwidth = NULL,
+    mu = NULL
+  )*
~~~

The maximum likelihood estimates with von Mises and wrapped Cauchy distributions or non-parametric kernel density estimates (`parametric=FALSE`) utilize the corresponding functions of the **circular** package (Agostinelli and Lund 2017). Our work was motivated by a need to fit circular distributions to animal movement data that exhibited peaks around two different directions (Hodel and Fieberg 2021). Therefore, we also provide the possibility to perform maximum likelihood estimation for a mixture of two von Mises distributions. The implementation for these mixtures is derived from the expectation-maximization algorithm of the **movMF** package (Hornik and Grün 2014) but with the added ability to fix and not optimize the two location parameters (using the `mu` argument to the `fit_angle()` function). Fixing the location parameters allows us to accommodate *a priori* assumptions specific to animal movement data (e.g., that there is no inherent bias for an animal to turn left or right). A similar function (`fit_steplength()`) is also available for linear random variables and a variety of common linear distributions.

The quantiles and CDF of the wrapped Cauchy distribution and the quantiles of the mixed von Mises distribution cannot be found analytically, and they are also not part of the **circular** package (Agostinelli and Lund 2017). Therefore, we implemented them numerically based on the wrapped G distribution of the **wrapped** package (Nadarajah and Zhang 2017) and spline interpolation, respectively (see Appendix A). Finally, we also provide PDF, CDF, random number generation, and quantiles for kernel density estimates with the functions `rdens(n, density)`, `ddens(n, density)`, `pdens(n, density)`, and `qdens(n, density)`. These functions all take as input a list containing information about the kernel density estimate, i.e., the output of

~~~
*R> density <- fit_angle(theta, parametric = FALSE)*
~~~

and similar for `fit_steplength()`.

## 4. Circular-linear copulae

In the following sections, we will introduce copulae that can be used to model bivariate distributions of circular-linear pairs of random variables, (Θ, *X*). Such copulae have as their domain not the unit square, but the surface of a cylinder of unit height and unit circumference (hence the name of the package, “**cylcop**”). This means that they must fulfill a boundary condition analogous to Equation 17

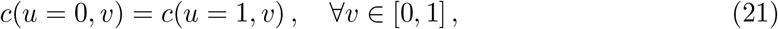

where we assumed that *u* is the dimension of the copula that will be transformed to the circular variable Θ. Since we are concerned with modeling movement data (i.e., joint turn angle and step length distributions), and animals usually do not have an inherent bias to turn left or right, we will focus on turn angle distributions that are symmetric around *θ* = 0. From Section 2.2, we know that this means that the copulae must not only fulfill Equation 21, but must also be symmetric in *u* around *u* = 0.5 (Equation 15)

With the **cylcop** package being an extension of the **copula** package (Hofert *et al*. 2020; Jun Yan 2007; Ivan Kojadinovic and Jun Yan 2010; Marius Hofert and Martin Mächler 2011), we follow their framework and implement the copulae we introduced above as “S4”-classes. For each of them, there are methods to calculate the PDF (`dcylcop()`), CDF (`pcylcop()`) and (inverse of) the conditional distribution (`ccylcop()`), as well as to generate random samples (`rcylcop()`).

### 4.1. von Mises copula

Johnson and Wehrly (Johnson and Wehrly 1978) present in section 3 of their paper an approach to obtain a circular-linear PDF with given marginal densities:

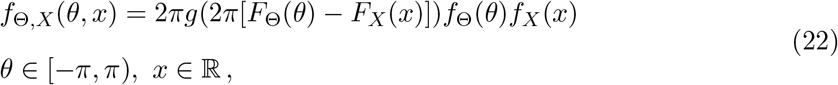

where *g* is some density on the circle. Even though it is not mentioned in that work, a comparison of Equations 22 and 3 reveals that 2*πg*(2*π*[*u* – *v*]) = *c*(*u, v*) must be a copula density. This has been recognized by many researchers, see e.g., García-Portugués *et al*. (2013), who chose as function *g* the von Mises density and applied this concept to the joint modeling of wind direction and concentration of SO_2_. We implement this copula as `cyl_vonmises()` in our **cylcop** package.

Next, we will derive the CDF of the von Mises copula and describe its implementation in the **cylcop** package. We will assume the von Mises density *g* to have as support the entire real line (see Equation 19), a mean *μ*_0_, and a concentration *κ*_0_. Substituting *q* = 2*πt* and 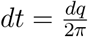 (in line 4, below), we can integrate the copula density

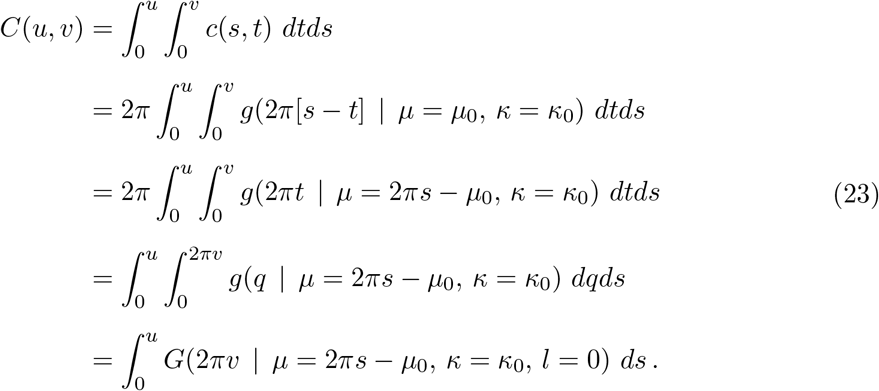

Going from the second to the third line, we used the symmetry properties of *g*, which can be seen in Figure 2. The innermost integral in the fourth line is just the von Mises CDF *G*(2*πv* | *l* = 0) with a lower limit of integration *l* = 0 (see Equation 20). Since there is no analytic solution, the final integral would have to be solved numerically. As mentioned in Section 3, the **circular** package implements the von Mises CDF as a periodic function (i.e., following Equation 20), and we could implement `pcylcop(cyl_vonmises(mu = mu_0, kappa = kappa_0))` by numerically integrating from *s* = 0 to *s* = *u* the function

~~~
*R> circular::pvonmises(q = (2 - .Machine$double.eps) * pi * v,
+    mu = 2 * pi * s - mu_0,
+    kappa = kappa_0,
+    from = 0
+  )*
~~~

Since *G* is periodic on the half-open interval [0, 2*π*), we multiply not with 2, but a number very close to 2. While this approach is illustrative, it is, however, computationally very slow. Therefore, the actual implementation of the CDF of the von Mises copula in the **cylcop** package approximates the double integral in Equation 23 with truncated infinite sums of Bessel functions. The function *g* can be expressed as an infinite series of Bessel functions *I_j_* of increasing order *j*

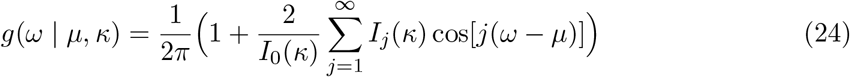

From this, we can arrive at the following expression for the bivariate CDF (the derivation can be found in Appendix B)

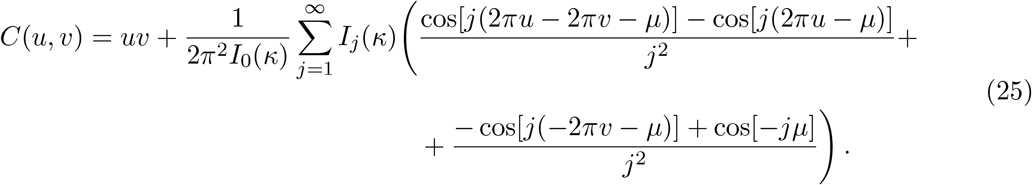

With density and distribution functions given above, what remains is the conditional distribution, which can be derived following Equation 23 as

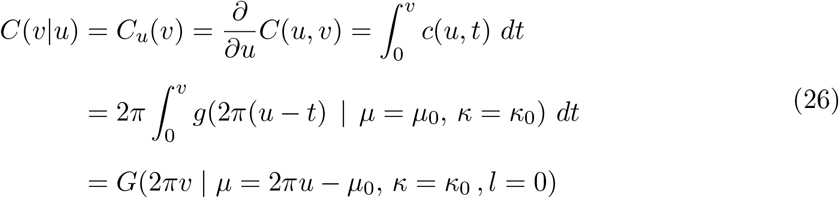

Its inverse (which we need for the generation of copula samples) is implemented as

~~~
*R> 1 / (2 * pi) * circular::qvonmises(p = v,
+    mu = 2 * pi * u - mu_0,
+    kappa = kappa_0,
+    from = 0
+  )*
~~~

#### Usage and properties

The von Mises copula (Figure 3) can be instantiated with

~~~
*R> cylcop::cyl_vonmises(mu = 0, kappa = 1, flip = FALSE)*
~~~

**Figure 3:**
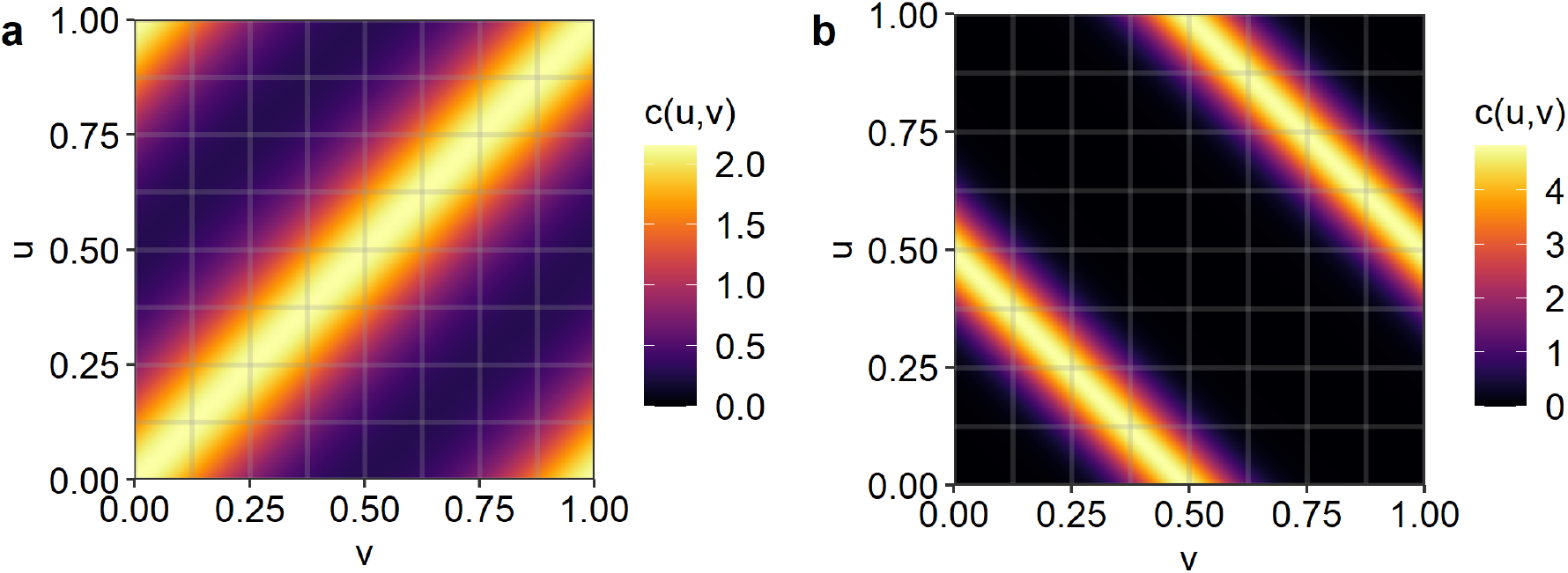
PDFs of `cyl_vonmises` copulae. Left: `mu = 0, kappa = 1`, right: `mu = pi, kappa = 4, flip = TRUE.`

The parameter `mu` shifts the PDF in the *v*-direction, `kappa` determines the concentration and `flip` indicates whether the copula should represent positive (`flip = FALSE`) or negative (`flip = TRUE`) correlation. With `kappa = 0`, we have the independence copula Π. With `kappa = Inf` and `flip = FALSE`, or `flip = TRUE`, we get the upper (*M*), or lower (*W*) Fréchet–Hoeffding bound, respectively. While the `cyl_vonmises` copula is indeed a circular-linear copula, and hence periodic, it is not symmetric, which limits its usefulness for the modeling of movement data. The copulae introduced in the following sections, on the other hand, have densities that are both periodic and symmetric in *u*.

### 4.2. Copulae with specific sections

The sections in *v* of a copula *C*(*u, v*) are defined as a set of functions, *C*(*u*_0_, *v*), from [0, 1] to [0, *u*_0_]: {*v* ↦ *C*(*u*_0_, *v*) | *u*_0_ ∈ [0, 1]}. Note that the upper bound of the range is *u*_0_ because *C*(*u*_0_, 1) = *u*_0_. The section in *v* at *u*_0_, *C*(*u*_0_, *v*), is the curve of intersection of the copula surface with the plane defined by *u* = *u*_0_, which is parallel to the *v*-axis. Furthermore, it is also proportional to the conditional distribution function *C*(*u*_0_, *v*) = *u*_0_ Pr[*V* ≤ *v* | *U* ≤ *u*_0_] (Nelsen, Quesada-Molina, and Rodríguez-Lallena 1997). Below, we show that specific copulae with quadratic or cubic sections in the linear direction *v* have periodic and symmetric densities in the circular direction *u*.

#### Quadratic sections

Following section 3.2.5 in Nelsen (2006), we note that copulae with quadratic sections in *v* must be of the form

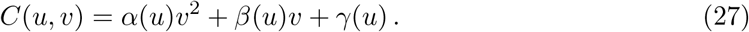

Since *C*(*u*, 0) = 0 = *γ*(*u*) and *C*(*u*, 1) = *u* = *α*(*u*) + *β*(*u*), we can define *ψ*(*u*) = –*α*(*u*) = *β*(*u*) – *u* to obtain

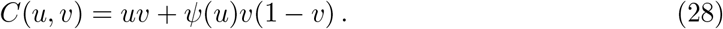

Due to the constraints that *C*(0, *v*) = 0 and *C* (1, *v*) = *v*, it is necessary that *ψ*(0) = *ψ*(1) = 0. Additionally, *ψ* must be chosen so that *C* is 2-increasing. In Corrolary 3.2.5, Nelsen (2006) states 3 necessary and sufficient conditions for this to be the case. We will now add a fourth one to make *c*(*u, v*), the copula density, periodic in *u*:

1. *ψ*(*u*) is absolutely continuous on [0, 1].
2. |*ψ*′(*u*)| ≤ 1 almost everywhere on [0, 1].
3. |*ψ*(*u*)| ≤ min(*u*, 1 – *u*) ∀*u* ∈ [0, 1].
4. *ψ*′(*u*) must be periodic on [0, 1].

The fourth condition is easily derived:

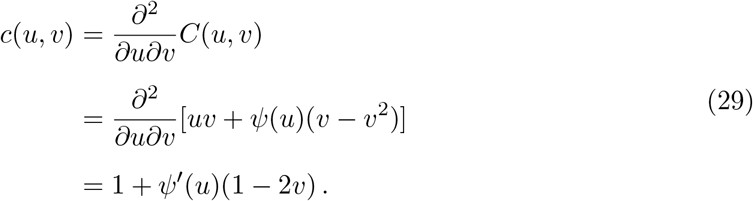

Since *ψ*′(*u*) is the only term containing *u*, requiring *c*(*u, v*) to be periodic in *u* means that *ψ*′(*u*) must be also periodic on [0, 1]. We will therefore choose

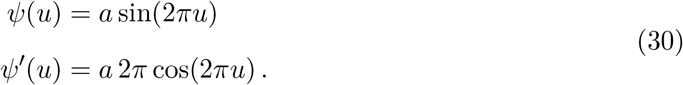

This function clearly satisfies *ψ*(0) = *ψ*(1) = 0 and fulfills property 1. It also satisfies properties 2 and 3 but only for certain values of *a*, – 1/2*π* ≤ *a* ≤ 1/2*π* (see Appendix C). Finally, note that *c*(*u, v*) is not only periodic in *u* on [0, 1], but also symmetric:

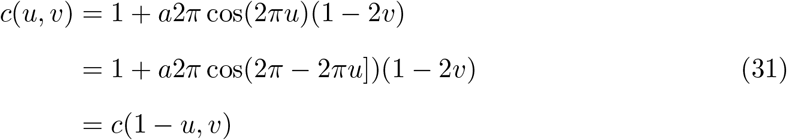

What remains to derive is the conditional distribution

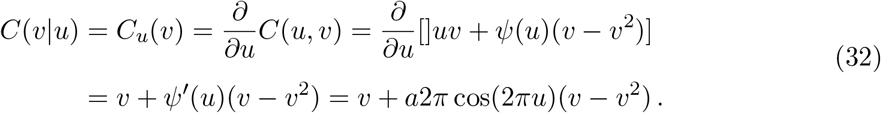

The inverse, which we need to generate random samples from the copula, is

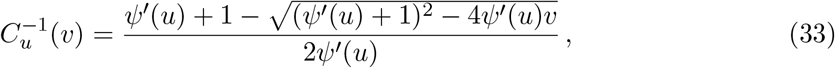

since

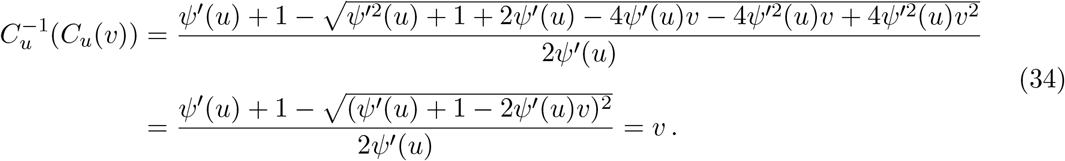

In summary, our circular-linear copulae with quadratic sections have the following properties:

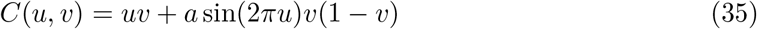

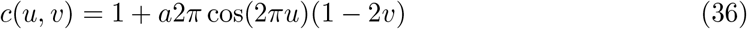

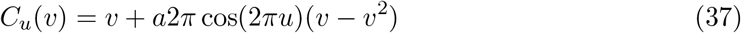

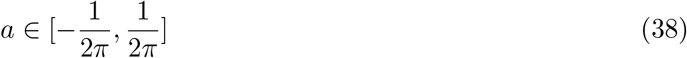

#### Cubic sections

Following an analogous path of reasoning about boundary conditions as in the quadratic case and according to equation 3.2.13 in Nelsen (2006), a copula with cubic sections must be of the form

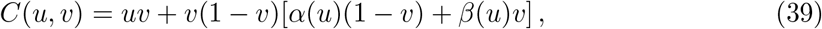

where *α*(0) = *α*(1) = *β*(0) = *β*(1) = 0. We will show now that with

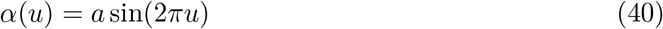

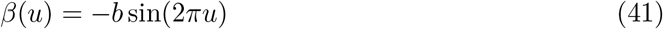

and certain values of *a* and *b*, *C* fulfills all requirements of a circular-linear copula with periodic density.

First, it is immediately clear that *α*(0) = *α*(1) = *β*(0) = *β*(1) = 0 holds for any values of *a* and *b*. Next, we need to show that our choice of *α* and *β* leads to a 2-increasing function *C*. To this end, we use theorem 3.2.8 in Nelsen (2006), which states that *α*(*u*) and *β*(*u*) must both be absolutely continuous, which is clearly satisfied. Furthermore, it is also necessary that the following set be non-empty for almost all *u* ∈ [0, 1]

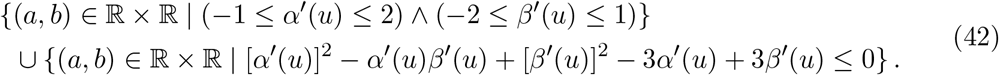

The parameter values in the first set of the union are the ones that fulfill

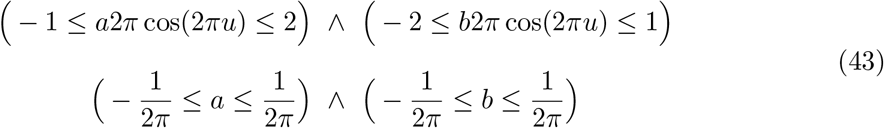

The second set of the union is somewhat more difficult to derive:

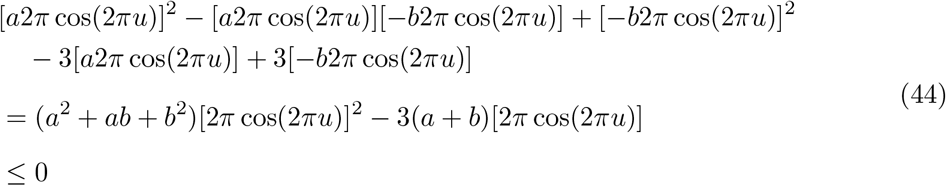

For given values of *a* and *b*, the difference is largest, when *u* = 0.5 and [2*π* cos(2*π*0.5)] = –2*π*, or when *u* = 1 and [2*π* cos(2*π*)] = 2*π*. Focusing on the first case:

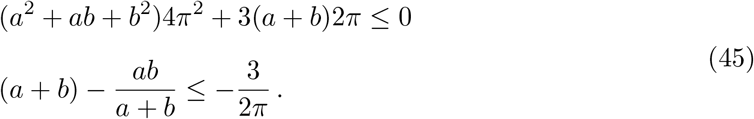

The first equation is a quadratic form and describes a closed set bounded by an ellipse. It is easy but tedious to show that its center point is at (*a* = –1/(2*π*), *b* = –1/(2*π*)), the angle between its minor axis and the “a-axis” is 0.25*π* and the distance between the center and the focal points is 1/*π*. The length of the semi-minor axis is 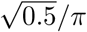. The second case (*u* = 1) leads to

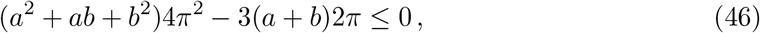

an ellipse that has its center point at (*a* = 1/(2*π*), *b* = 1/(2*π*)). The angle between its minor axis and the “*a*-axis” is 0.25*π*, the distance between the center and the focal points is 1/*π*, and the length of the semi-minor axis is 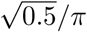. From the lengths and directions of their semi-minor axes and their center positions, it is clear that the two ellipses only touch at one point (*a* = 0, *b* = 0). Finally, the set of admissible parameter values is the union between the closed set bounded by the square described in Equation 43 and this set containing only (*a* = 0, *b* = 0), which is a subset of the first one (see Appendix D). What remains to show is that *c*(*u, v*) is periodic (and symmetric) in *u*.

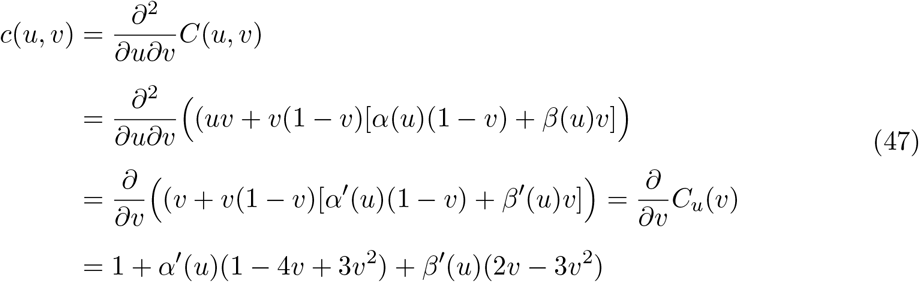

Since for *α*′(*u*) = *a*2*π* cos(2*πu*) and *β*′(*u*) = *b*2*π* cos(2*πu*) it holds that *α*′(*u*) = *α*′(1 – *u*) and *β*′(*u*) = *β*′(1 – *u*), the copula density is not only periodic in u, *c*(0, *v*) = *c*(1, *v*), but also symmetric, *c*(*u, v*) = *c*(1 – *u, v*). The inverse of the conditional distribution (line 3 of the equation above) is implemented analytically in the **cylcop** package. The resulting expression is however very unwieldy and is therefore omitted here.

#### Usage and properties

Quadratic and cubic sections copulae (Figure 4) can be generated using

~~~
*R> cylcop::cyl_quadsec(a = 1 / (2 * pi))
R> cylcop::cyl_cubsec(a = 1 / (2 * pi), b = 1 / (2 * pi))*
~~~

**Figure 4:**
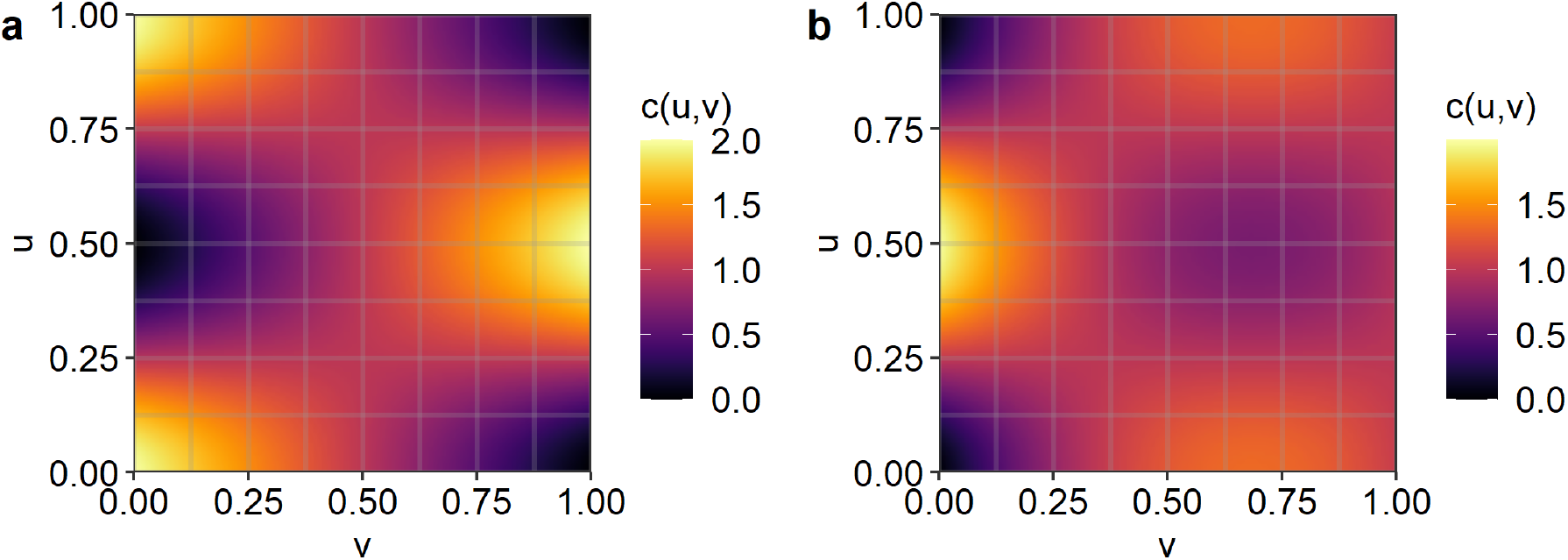
PDFs of copulae with specific sections in *v*. Left: `cyl_quadsec(a = 1/(2*pi))`, right: `cyl_cubsec(a = −1/(2*pi), b = 0.01)`.

The parameters a and b can take values from [–1/(2*π*), 1/(2*π*)] (see Equation 43) and with `a = 0` (and `b = 0`) we obtain the independence copula Π. The Fréchet–Hoeffding bounds can, however, not be attained. With *a* > 0 (and *b* < 0) the copula captures positive correlation between *u* and *v* for *u* ∈ [0, 0.5] and negative correlation for *u* ∈ [0.5, 1]. The opposite is true for *a* < 0 (and *b* > 0).

### 4.3. Combinations of copulae

#### Rotated copulae

We have seen in Section 2.2 that orthogonal reflections or “rotations” of copula densities are themselves copula densities. Furthermore, any convex linear combination of copulae is a copula (Nelsen 2006). We will now show that taking the arithmetic mean between any linear-linear copula *C*(*u, v*) and its orthogonal reflection along the line u=0.5, *C*_0.5*π*_, produces a circular-linear copula

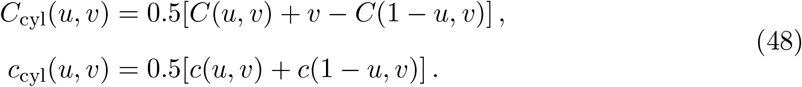

This copula density is not only periodic, but also symmetric in *u*:

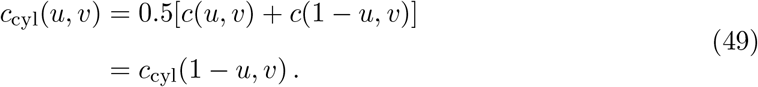

We can also “periodically shift” the density by 0.5 in *u*-direction, i.e., *c*_cyl_((*u* + 0.5) mod 1, *v*) or

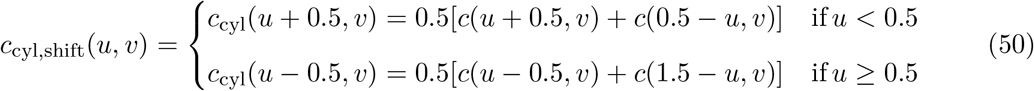

It can be shown with simple geometric arguments that the distribution of such a shifted copula is

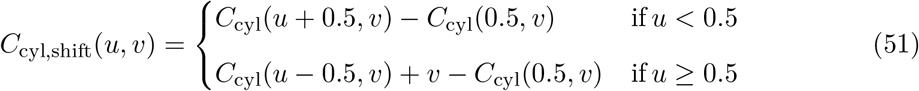

That this is indeed a copula can be easily proven and *C*_cyl,shift_ is in fact a special case of a shuffle of *C*_cyl_ see, e.g., section 3.2.3 of Nelsen (2006) or section 3.6 of Durante and Sempi (2015).

#### Rectangular patchworks of copulae

Another method of constructing copulae is from rectangular patchworks (Durante, Saminger-Platz, and Sarkoci 2009). The unit square, i.e., the domain of the copula, is divided into rectangular regions *R_i_* that can only potentially share points at their boundaries. For points (*u, v*) outside those rectangles, the copula *C*(*u*, *v*) is equal to some copula *C*_bg_(*u, v*) (“bg” for background). For points that are an element of rectangle *R_i_*, the copula is equivalent to a 2-increasing function *F_i_(*u*, *v**) : *R_i_* → [0, 1] obtained by some transformation of a copula *C_i_*(*u*, *v*) so that *F_i_* = *C*_bg_ at the boundaries of *R_i_*. By choosing appropriate rectangles and copulae *C_i_* and *C*_bg_, we can construct circular-linear copulae.

Specifically, we define 2 rectangles that are mirror images of each other with respect to the line *u* = 0.5: *R*_1_ = [*u*_1_, *u*_2_] × [0, 1] and *R*_2_ = [1 – *u*_2_, 1 – *u*_1_] × [0, 1] with 0 ≤ *u*_1_ < *u*_2_ ≤ 0.5. The patchwork-copula is then periodic and symmetric in *u* if the following two conditions are satisfied

- *C*_2_ is the orthogonal reflection of *C*_1_ with respect to the line *u* = 0.5, i.e., *C*_2_(*u*, *v*) = *C*_1, 0.5*π*_ (*u, v*) = *v* – *C*_1_(1 – *u, v*).
- *C*_bg_ is a copula that is periodic and symmetric in *u*, or *u*_1_ = 0 and *u*_2_ = 0.5, i.e., the two rectangle cover the entire unit square.

The explicit distribution of a general patchwork-copula (of which the above-described copula is a special case) is given in Theorem 2.2 in Durante *et al*. (2009) and is repeated here for our specific two rectangles:

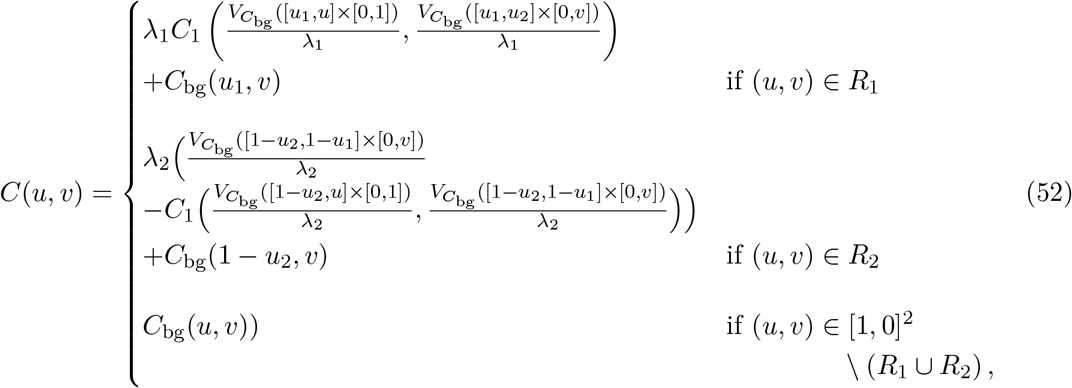

with λ*_i_* = *V*_*C*_bg__ (*R_i_*) (see Equation 1 for the definition of the C-volume). In case the two rectangles cover the entire unit square and *C*_bg_(*u, v*) = *uv* is the independence copula, this equation drastically simplifies to

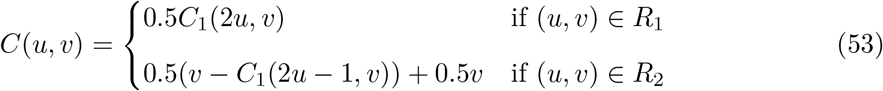

since in this case λ*_i_* = 0.5, and *V*_*C*_indep__([0, *u*] × [0, *v*]) = *uv*.

The copula PDF is obtained by taking the derivatives of the CDF (see Appendix E for a detailed derivation), which results in

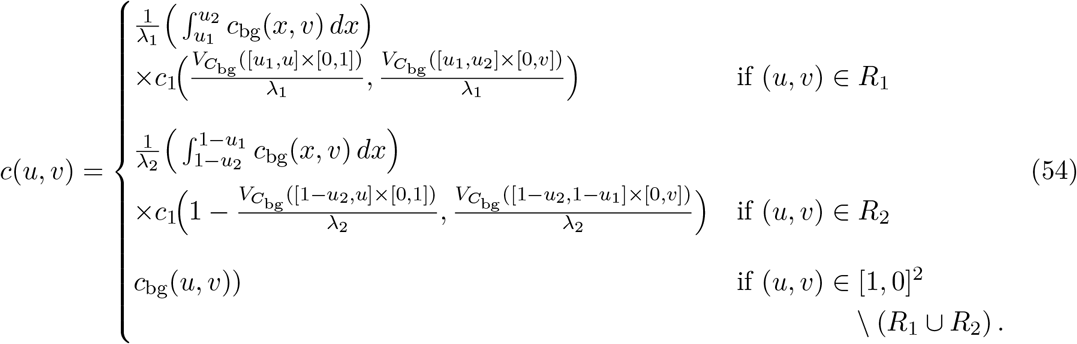

As was the case for the distribution, this equation is a lot simpler when the two rectangles cover the entire unit square and *C*_bg_(*u*, *v*) = *uv*:

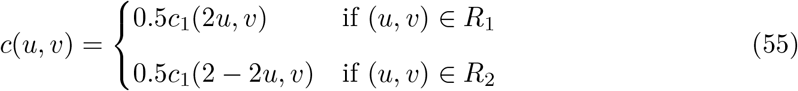

To sample from the patchwork-copula, we follow the algorithm described in section 2 of Durante *et al*. (2009): Generate a sample (*u*, *v*) from *C*_bg_(*u*, *v*). If this point does not fall within any rectangle, take it as a sample from *C*. If it falls within rectangle *i*, generate a sample (*x_i_, y_i_*) from *C_i_* and transform it to 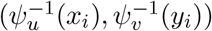, where 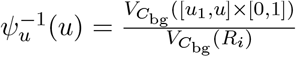 and 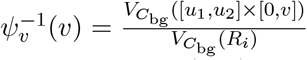. In the special case where the two rectangles cover the entire unit square and *C*_bg_(*u*, *v*) = *uv*, the transformation simplifies to (*x*_1_ /2, *y*_1_) for *R*_1_ and ((*x*_2_ + 1)/2, *y*_2_) for *R*_2_.

With the algorithm described above, it is not necessary to find the inverse of the conditional distribution of the patchwork-copula to generate samples from it. Nevertheless, we implemented the conditional distribution and its inverse using numeric integration of the copula density and numeric inversion.

#### Usage and properties

A circular-linear copula consisting of a linear combination of a linear-linear copula and its “90 degrees rotation” can be obtained as

~~~
*R> cyl_rot_combine(copula, shift = FALSE)*
~~~

where `copula` must be a `copula` object from the **copula** package (Hofert *et al*. 2020; Jun Yan 2007; Ivan Kojadinovic and Jun Yan 2010; Marius Hofert and Martin Mächler 2011). This gives a copula with a correlation structure that leads to an X-shaped PDF. With `shift = TRUE`, the same copula, but with its PDF “periodically shifted” by 0.5 in *u*-direction, is obtained, which has a diamond-shaped density. This is illustrated in Figures 5a and 5b with the following copulae

~~~
*R> cop_lin <- copula::rotCopula(claytonCopula(8), flip = TRUE)
R> cop_a <- cyl_rot_combine(cop_lin, shift = FALSE)
R> cop_b <- cyl_rot_combine(cop_lin, shift = TRUE)*
~~~

**Figure 5:**
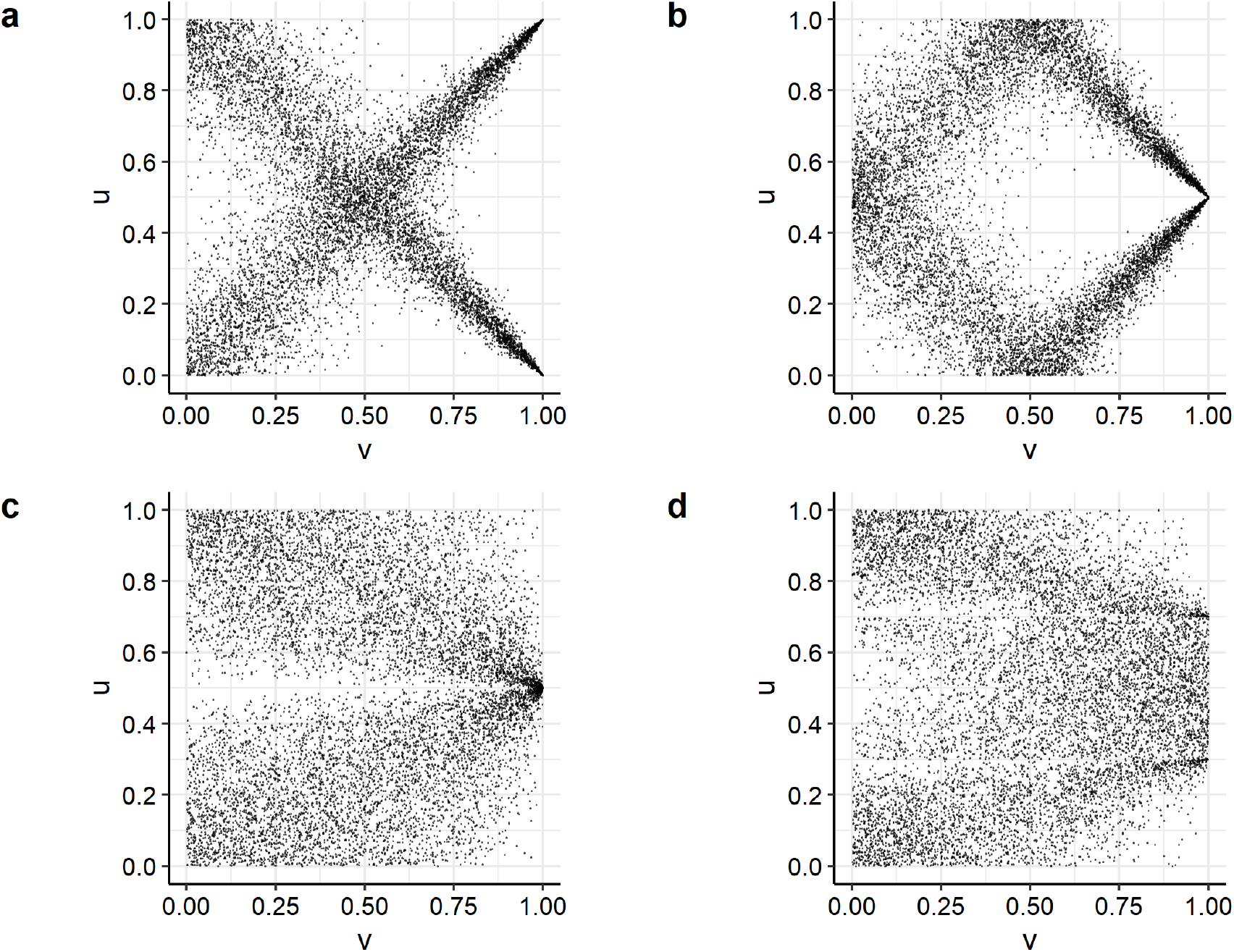
Scatterplots of 10,000 samples drawn from the following copulae (see text). a: `cop_a`, b: `cop_b`, c: `cop_c`, d: `cop_d`.

A circular-linear copula generated from a patchwork of copulae can be obtained as

~~~
*R> cyl_rect_combine(copula,
+    background = indepCopula(),
+    low_rect = c(0, 0.5),
+    up_rect = “symmetric”,
+    flip_up = TRUE
+  )*
~~~

where `copula` is the copula on which the functions in the two rectangles are based and must be a copula-object from the **copula** package (Hofert *et al*. 2020; Jun Yan 2007; Ivan Kojadinovic and Jun Yan 2010; Marius Hofert and Martin Mächler 2011), or a `cyl_vonmises` object, whereas the copula outside the two rectangle (`background`) can be any circular-linear or linear-linear copula. The direction of correlation can be tuned by “rotating” the copulae on which the functions in the rectangles are based using `flip_up` and by changing the correlation structure of the background-copula. This is illustrated in Figures 5c and 5d with the following copulae

~~~
*R> cop_lin <- copula::rotCopula(claytonCopula(1.1), flip = TRUE)
R> cop_c <- cyl_rect_combine(cop_lin)
R> cop_d <- cyl_rect_combine(cop_lin,
+    background = cyl_quadsec(1 / (2 * pi)),
+    low_rect = c(0, 0.3),
+    up_rect = “symmetric”
+  )*
~~~

## 5. Dependence and parameter estimates

### 5.1. Parameter estimation

Given a set of *n* i.i.d. observations of a pair of circular and linear variables (*θ_i_, x_i_*), an assumed copula family, 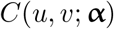, and assumed marginal distributions, 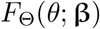 and 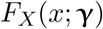, we could, in principle, estimate the parameter vector 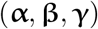 by maximizing the log-likelihood based on the joint PDF of Equation 3. However, this approach would require optimizing a large number of parameters simultaneously, and mis-specification of the marginal distributions would lead to biased estimators of the copula parameters and vice versa.

If we knew the true marginal distributions, we could transform our data to observations from the copula and maximize the log-likelihood

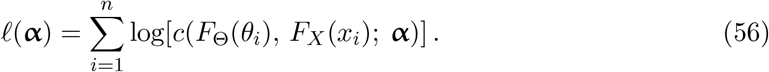

This leads to two alternative approaches for estimating copulae parameters, in which estimates of *F*_Θ_ and *F_X_* are plugged into Equation 56. These estimates, 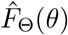 and 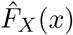, can either be obtained parametrically via maximum likelihood (Joe and Xu 1996; Joe 1997, 2005), or non-parametrically as pseudo-observations, i.e., draws from the empirical copula (Section 2.1). Note that in the parametric case, misspecified marginal distributions still lead to biased estimators of copula parameters (Fermanian and Scaillet 2005). The number of parameters that have to be simultaneously optimized is, however, reduced in both cases. When using estimates, 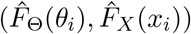, instead of the true marginal distributions in Equation 56, 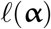 is no longer a true log-likelihood. Further, when using the empirical copula (Equation 7), observations are also no longer i.i.d. since the non-parametric estimates of the marginal distributions clearly depend on the entire data set (Hofert *et al*. 2018). The estimation of copula parameters using non-parametrically estimated marginals is referred to as maximum pseudo-likelihood estimation (MPLE, see Oakes 1994; Genest, Ghoudi, and Rivest 1995; Shih and Louis 1995; Tsukahara 2005) and is implemented in the **cylcop** package as

~~~
*R> optML(copula, theta, x, parameters, start)*
~~~

The arguments of this function, which are based on `copula::fit_copula()` from the **copula** package, are a circular-linear copula object (`copula`), the circular (`theta`) and linear (`x`) components of the measurements, the parameters of the copula that are to be optimized (`parameters`), and their initial values for that optimization (`start`).

For many linear-linear copulae, parameter estimates can be derived analytically from estimates of a correlation measure, such as Spearman’s rho or Kendall’s tau (Oakes 1982; Genest 1987; Genest and Rivest 1993); analytical solutions are not possible, however, for most of the copulae introduced in this paper, and we, therefore, had to resort to using numerical optimization methods. The exception are patchwork copulae, where the rectangles span the entire unit square. Their parameters can be estimated using Kendall’s tau implemented with the function

~~~
*R> optCor(copula, theta, x, method = “tau”)*
~~~

For all other circular-linear copulae, we can carry out a grid-search to determine the copula parameters that minimize the difference in a correlation measure (Section 5.2) between the empirical and parametric copula. This approach assumes the absolute values of copula parameters increase monotonously with increasing circular-linear correlation and is implemented as

~~~
*R> optCor(copula, theta, x, method = c(“cor_cyl”, “mi_cyl”))*
~~~

The two methods `“cor_cyl”` and `“mi_cyl”` refer to a non-parametric estimate of a circular-linear correlation coefficient and mutual information, respectively (see Section 5.2). The parameter values obtained in this way serve as reasonable starting values for MPLE, the choice of which is especially important when the parameter hypersurface is rugged and highdimensional.

Finally, model selection can be performed according to Akaike’s information criterion, which is a common (e.g., Chen and Fan 2005; McNeil, Frey, and Embrechts 2015; Joe 2014), but for MPLE not uncontroversial approach (see Grønneberg and Hjort 2014, but also Jordanger and Tjøstheim 2014).

### 5.2. Measures of correlation

Parametric measures of circular-linear correlation require normally distributed linear random variables (Johnson and Wehrly 1977; Mardia 1976) and are therefore not generally applicable to all circular-linear data. Instead, we used a non-parametric estimate of a circular-linear correlation coefficient, *D*, that was introduced by Mardia (1976) and applied by Solow, Bullister, and Nevison (1988) or, more recently, Tu (2015). It is estimated by ranking the data, i.e., from the pseudo-observations

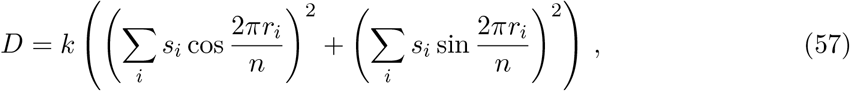

where *k* is a normalization constant, *r_i_* is the circular (with –*π* as reference) and *s_i_* the linear rank of observation *i*. However, this coefficient can only take values between 0 and 1 and thus cannot distinguish between positive and negative correlation.

We also implemented a correlation measure based on mutual information, i.e., the Kullback-Leibler divergence of the product of the marginal distributions from the joint distribution (see e.g., Leguey, Larrañaga, Bielza, and Kato 2019).

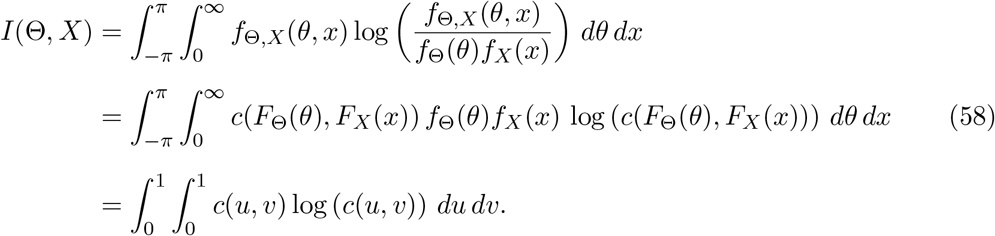

The last equality was obtained by setting *u* = *F*_Θ_(*θ*) and *v* = *F_X_* (*x*). From this expression, it is immediately clear that the mutual information is the negative of the differential entropy associated with *c*(*u, v*) (Ma and Sun 2011; Calsaverini, Vicente, Systems, and Artes 2009). If we could easily carry out the integration in the last line of Equation 58, we could directly determine the parameter values of the chosen copula family that give the correct mutual information. Unfortunately, the integral is computationally difficult to calculate in most cases. However, Tenzer and Elidan (2016) have demonstrated that the mutual information is monotonous in the dependence parameter for a wide range of bivariate copulae. The rough optimization framework described above can therefore be expected to work well with mutual information as a criterion for dependence. We can obtain an estimate of the mutual information by binning the data (measurements of the circular and the linear random variable) and taking the average of the point-wise mutual information (based on the frequency of data points in the bins) over all bin combinations. Note also that the mutual information of the data is equal to the mutual information of the empirical copula. If required, the mutual information can be normalized to an interval [0, 1] (zero meaning independence copula and 1 meaning upper Fréchet-Hoeffding copula) with the entropy *H*:

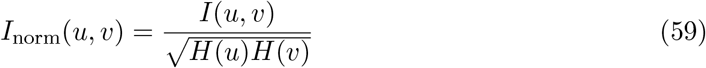

Finally, if we take it as fact that the true copula is symmetric in *u*, we can symmetrize the empirical copula (set the values of *u* that are larger than 0.5 to 1 – *u*). The two correlation measures are implemented as

~~~
*R> cor_cyl(theta, x)
R> mi_cyl(theta, x, normalize = TRUE, symmetrize = FALSE)*
~~~

## 6. Coded examples

To begin with, we will load the necessary packages, reduce the verbosity of the package, and generate two `cylcop` objects with which we will demonstrate the functionality of the **cylcop** package. First, we generate a copula with cubic sections

~~~
*R> library(“copula”)
R> library(“cylcop”)
R> cylcop_set_option(silent = TRUE)
R> copula_1 <- cyl_cubsec(a = 0.05, b = −1 / (2 * pi))*
~~~

Secondly, to demonstrate the level of complexity in the correlation structure that can be captured, we generate a copula consisting of a rectangular patchwork with rectangles [0.1, 0] × [0.4, 1] and [0.6, 0] × [0.9, 1]. Inside of the rectangles, the copula is derived from a Gumbel copula rotated by 180 degrees (first rectangle) and a Gumbel copula rotated by 180 and 90, i.e., 270, degrees. Outside the rectangles, the copula consists of the linear combination of a Frank copula and its 90-degree rotation.

~~~
*R> cop_lin <- copula::rotCopula(gumbelCopula(1.2), flip = TRUE)
R> cop_bg <- cyl_rot_combine(frankCopula(10), shift = FALSE)
R> copula_2 <- cyl_rect_combine(copula = cop_lin,
+    background = cop_bg,
+    low_rect = c(0.1, 0.4),
+    up_rect = c(0.6, 0.9),
+    flip_up = TRUE
+  )*
~~~

The densities of these two copulae are shown in Figure 6 and were produced using **cylcop**’s `cop_plot()` function

~~~
*R> cop_plot(copula_1, type = “pdf”, plot_type = “ggplot”, resolution = 200)*
~~~

**Figure 6:**
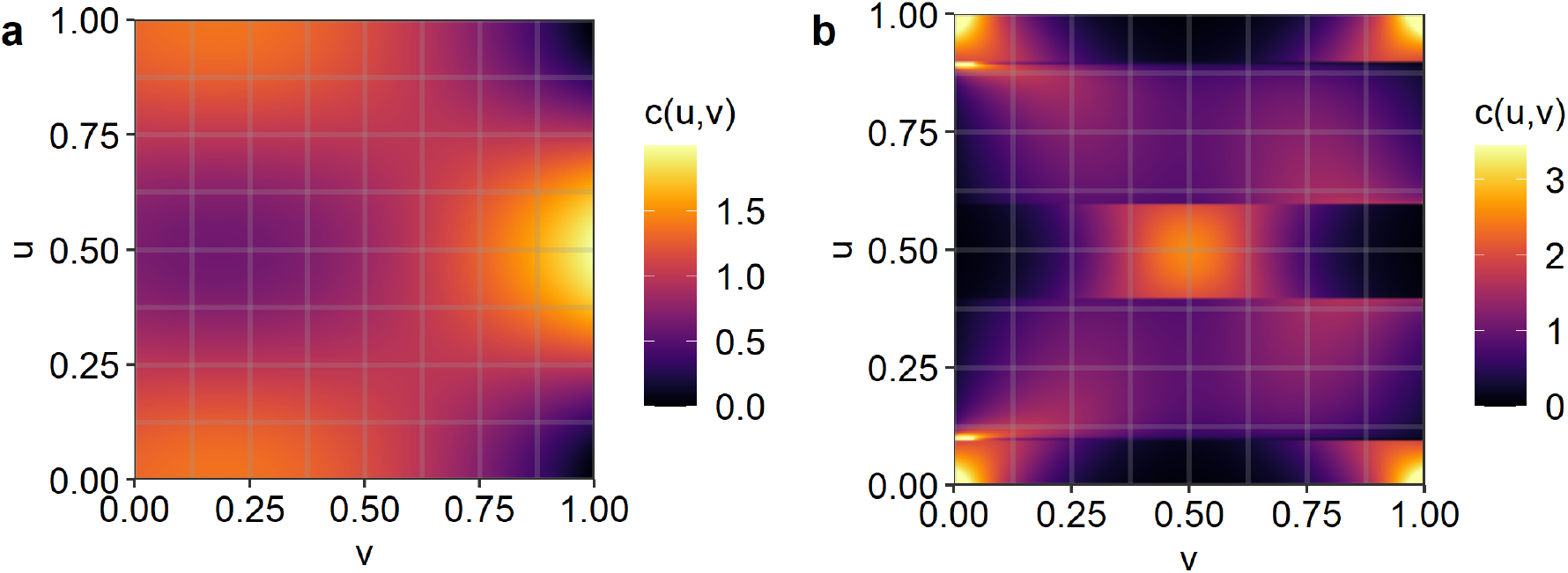
PDFs of a: cubic sections copula `copula_1` and b: rectangular patchwork copula `copula_2`.

Similarly, we could plot the CDF by setting `type = “cdf”`, and we can also plot 3-dimensional surfaces using the **rgl** package (Murdoch and Adler (2021); `plot_type = “rgl”`) or the **plotly** package (Sievert (2020); `plot_type = “plotly”`).

As we mentioned in the introduction, our motivation for deriving new copulae for correlated circular-linear data and for developing software for their implementation stems from a desire to model animal movement in discrete time (Hodel and Fieberg 2021). Thus, we also provide functions for simulating movement trajectories using copulae in the package. Selecting the first copula (`copula_1`), we can now generate a trajectory from 10,000 step lengths (distances between consecutive locations) and turn angles (changes in direction between successive locations) drawn from a joint distribution with a correlation structure described by that copula. As marginal distributions, we chose a gamma distribution for the step lengths and a mixed von Mises distribution for the turn angles.

~~~
*R> traj <- make_traj(n = 10000,
+    copula = copula_1,
+    marginal_circ = “mixedvonmises”,
+    parameter_circ = list(mu1 = 0, mu2 = pi, kappa1 = 2,
+       kappa2 = 1, prop = 0.7),
+    marginal_lin = “gamma”,
+    parameter_lin = list(shape = 3)
+  )*
~~~

Any distribution for which a quantile function is available can be used when specifying the linear marginal distribution (using argument `marginal_lin`). For the circular marginal distribution, the von Mises, the wrapped Cauchy, and a mixture of 2 von Mises distributions are available.

The output of `make_traj()` is a `data.frame` that contains the draws from the copula, the step lengths and turn angles, i.e., the draws from the copula transformed by the inverse CDFs of the marginal distributions, and the positions in space the animal would visit by moving according to those step lengths and turn angles. We have implemented several functions, below, to visualize simulated trajectories (see Figure 7).

~~~
*R> scat_plot(traj, periodic = TRUE)
R> traj_plot(traj)
R> cop_scat_plot(traj)
R> circ_plot(traj)*
~~~

**Figure 7:**
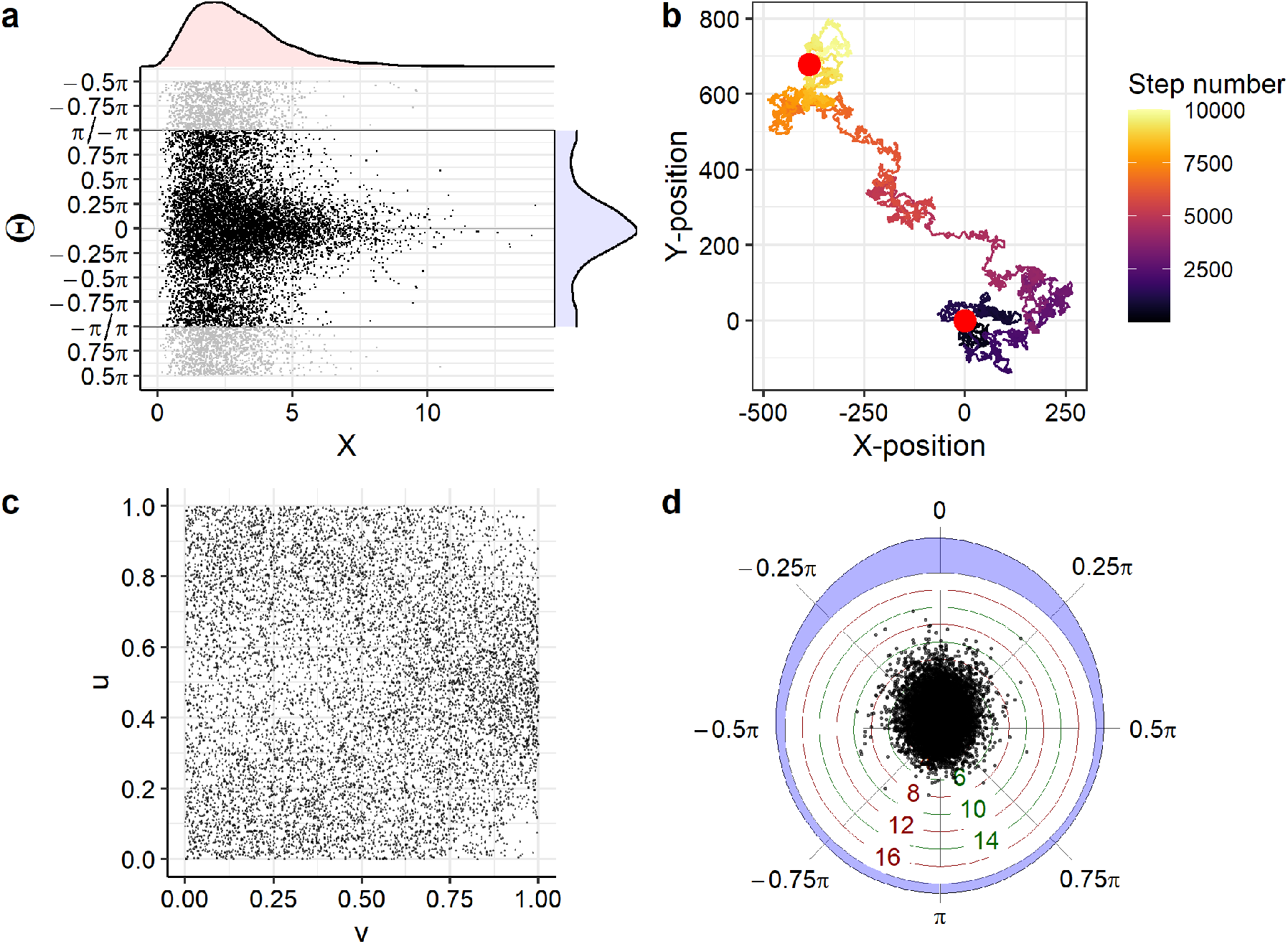
Circular-linear data representing turn angles and step lengths were simulated using `copula_1`, a copula with cubic sections, and marginal distributions given by gamma and mixed von Mises distributions. a: scatterplot of turn angles (Θ) and step lengths (*X*) with nonparametric density estimates in blue and red, respectively, obtained with `scat_plot()`. b: Plot of the trajectory derived from the sequence of turn angles and step lengths obtained with `traj_plot()`. c: scatterplot of the draws from `copula_1` obtained with `cop_scat_plot()`. d: circular scatterplot of turn angles and step lengths obtained with `circ_plot()`.

The argument `periodic` determines whether the scatterplot is periodically extended past the angles –*π* and *π*.

We provide several options for quantifying the correlation between circular and linear variables, which we demonstrate with our simulated data. The first available option is the circular-linear correlation coefficient described in Section 5.2.

~~~
*R> cor_cyl(theta = traj$angle, x = traj$steplength)*
[1] 0.07490922
~~~

The second option is to calculate the mutual information between the step lengths and turn angles. The mutual information can be normalized to lie in [0, 1] by dividing by the square root of the product of the entropies of the two random variables (see Equation 59). By specifying `symmetrize = TRUE`, we set the values of *u* of the empirical copula that are larger than 0.5 to 1 – *u*, ensuring that a symmetric circular-linear copula with perfect correlation has a normalized mutual information of 1 (see Appendix F)

~~~
*R> mi_cyl(theta = traj$angle,
+    x = traj$steplength,
+    normalize = TRUE,
+    symmetrize = TRUE
+  )*
[1] 0.02147514
~~~

As described in Section 5.2, neither correlation measure depends on the marginal distributions, which means that we obtain the same output, whether we take step lengths and turn angles as input, or the corresponding untransformed draws from the copula.

To facilitate model selection, we have written a function, `opt_auto()`, to autonomously fit 15 different circular-linear copulae (see Appendix G) to the data. The function returns a list containing descriptions of the copulae, `cylcop` objects, and AIC values associated with the fitted copulae.

~~~
*R> guess_fit <- opt_auto(theta = traj$angle, x = traj$steplength)*
~~~

The copula with the lowest AIC (−912.91) is a cubic sections copula with parameters *a* = 0.051 and *b* = –0.16, i.e., the copula with which we have generated the data (`copula_1`). The copulae with the second and third lowest AIC are a quadratic sections copula (*a* = 0.11, AIC = – 758.50, see Figure 8a) and a rectangular patchwork copula with symmetric rectangles spanning the entire unit square and the copula inside the rectangles based on a Frank copula with parameter *α* = 1.67 (AIC = –747.01, see Figure 8b).

**Figure 8:**
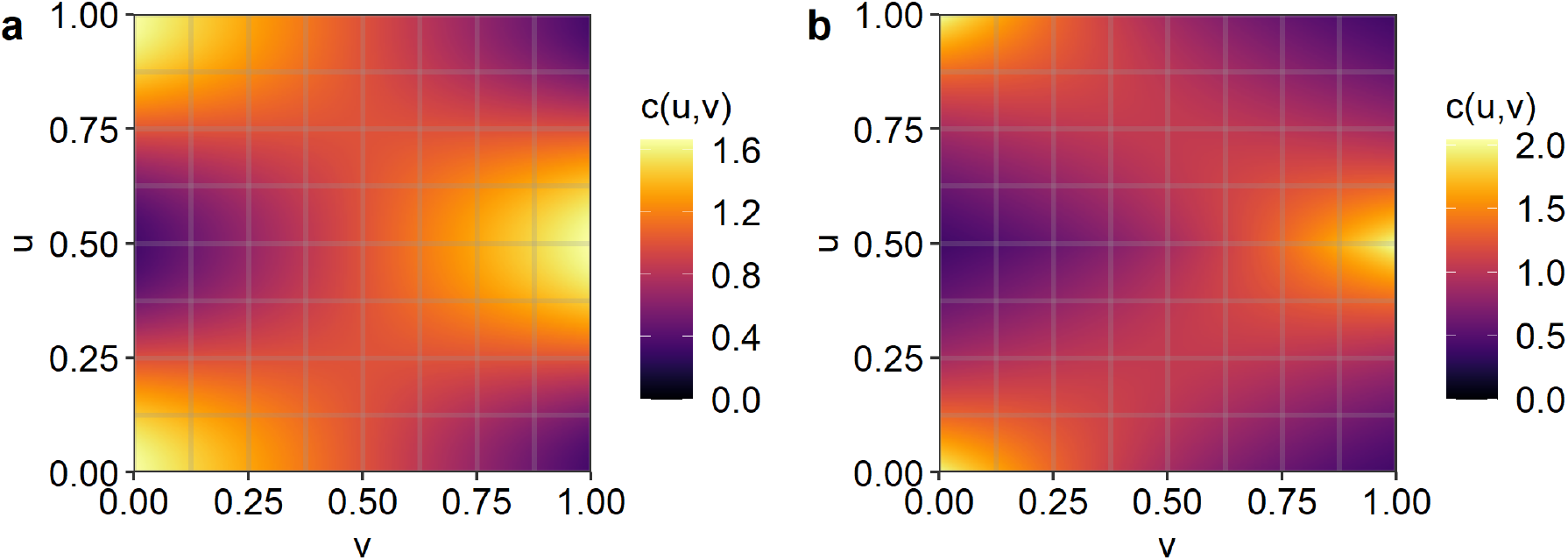
a: PDF of a quadratic sections copula with *a* = 0.11. b: PDF of a rectangular patchwork copula with symmetric rectangles spanning the entire unit square and the copula inside the rectangles based on a Frank copula with parameter *α* = 1.67.

**Figure 9:**
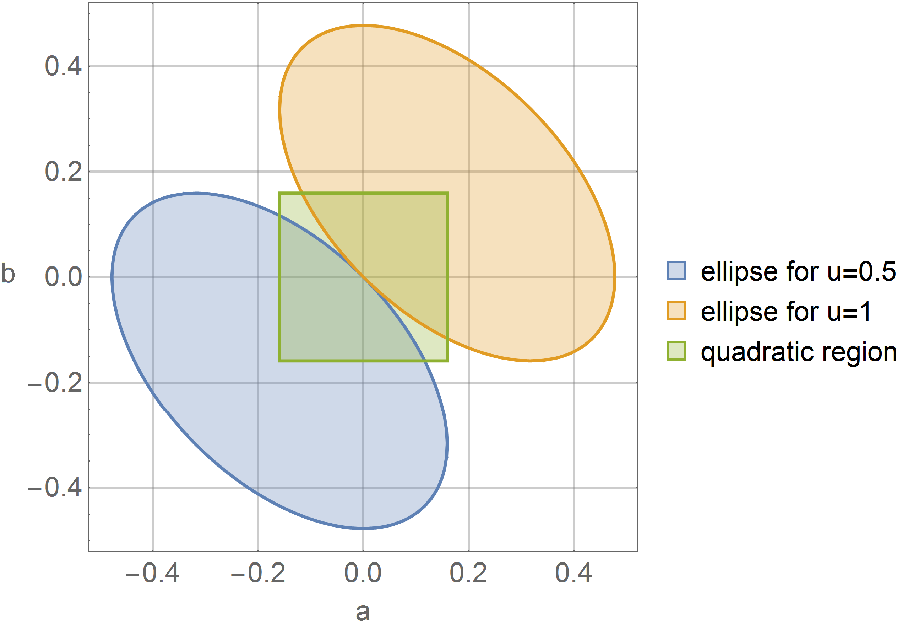
The set of permissible values of parameters *a* and *b* is the union of the square and the intersection of the ellipses (i.e. the square itself).

To fit a chosen copula to the data “by hand” we can first calculate starting values for MPLE using a correlation measure. `optCor()` first calculates the circular-linear correlation coefficient or the mutual information (specified by the argument `method`). Next, `n` samples are drawn from `copula` and the corresponding correlation measure is calculated. The parameters of the copula are changed and the process is repeated until the copula parameters giving the lowest difference between the correlation of the data and the correlation of the sample drawn from that copula are found with an accuracy specified by the argument `acc`. The parameter search is implemented as an exponential followed by a binary search.

~~~
*R> start_val <- optCor(copula = cyl_cubsec(0, 0),
+    theta = traj$angle,
+    x = traj$steplength,
+    acc = 0.01,
+    n = 1000,
+    method = “cor_cyl”,
+    parameter = “both”
+  )
R> start_val*
[1] 0.09084506 −0.07915494
~~~

Finally, copula parameters can be fit to the data via MPLE with the output of `optCor()` as starting value. The function `optML()` is based on `copula::fitCopula()` and returns a list containing the `cylcop` object with the optimized parameters, the maximum likelihood value and the corresponding AIC.

~~~
*R> optimized <- optML(copula = cyl_cubsec(),
+    theta = traj$angle,
+    x=traj$steplength,
+    parameters = c(“a”,”b”),
+    start = start_val,
+    lower = c(−0.1, −1 / (2 * pi)),
+    traceOpt = TRUE,
+    optim.method = “L-BFGS-B”,
+    optim.control = list(maxit = 100)
+  )
R> optimized*
$copula
Cub. sect. copula
a = 0.05103531
b = −0.1591549
$logL
[1] 458.4573
$AIC
[1] −912.9146
~~~

We also provide functions to fit the parameters of linear (`fit_steplength()`) and circular (`fit_angle()`) marginal distributions to the data using maximum likelihood. The **circular** (Agostinelli and Lund 2017) and **MASS** packages (Venables and Ripley 2002) provide excellent methods and functions for fitting almost any circular or linear distribution; our functions are basically just wrappers to these functions with some simplifications for easier usage. One exception is the mixed von Mises distribution, for which there was no method available to obtain maximum likelihood estimates in the **circular** package. We based our implementation on code from the **movMF** package (Hornik and Grün 2014) with the added feature of optionally fixing the mean directions.

~~~
*R> angle_distr <- fit_angle(theta = traj$angle,
+    parametric = “mixedvonmises”,
+    mu = c(0, pi)
+  )
R> unlist(angle_distr$coef)*
      mu1       mu2    kappa1    kappa2      prop
0.0000000 3.1415927 1.9634596 0.9549062 0.7108820
~~~

For both linear and circular marginal distributions, we can also obtain non-parameteric density estimates by setting `parametric = FALSE`, and passing a bandwidth (in this case, obtained using the `opt_circ_bw` function)

~~~
*R> bw <- opt_circ_bw(theta = traj$angle,
+    loss = “adhoc”,
+    kappa.est = “trigmoments”
+  )
R> angle_non_param <- fit_angle(theta = traj$angle,
+    parametric = FALSE,
+    bandwidth = bw
+  )
R> bw*
[1] 41.9904
~~~

Further details can be found in Agostinelli and Lund (2017). As is also described there, an ad-hoc estimation of the bandwidth used for the non-parametric density estimate can give wrong results, especially with multimodal distributions, and we compare density estimates obtained with different bandwidths in Appendix H). Finally, parametric and non-parametric density estimates can be used to simulate trajectories

~~~
*R> step_distr <- fit_steplength(x = traj$steplength, parametric = “gamma”)
R> unlist(step_distr$coef)*
    shape      rate
2.9803921 0.9965143
*R> traj_fit <- make_traj(n = 10000,
+    copula = optimized$copula,
+    marginal_circ = “dens”,
+    parameter_circ = angle_non_param,
+    marginal_lin = “gamma”,
+    parameter_lin = step_distr$coef
+  )*
~~~

## 7. Conclusions

We derived new copulae and implemented them in our **cylcop** package to facilitate modeling correlated circular-linear data where the circular variable is expected to be symmetric. We aimed to provide a thorough derivation of our circular-linear copulae together with a package that is easy to use, even as a black box. This should, on the one hand, allow application-oriented researchers to easily and quickly use these copulae to visualize and model their data and, on the other hand, provide statisticians and modelers with a starting point to further develop new methods.

In the future, we hope to integrate non-parametric copulae obtained from smoothing the empirical copula using e.g., Bernstein polynomials (Sancetta and Satchell (2004); Janssen, Swanepoel, and Veraverbeke (2012); Carnicero, Ausín, and Wiper (2013); García-Portugués *et al*. (2013); García-Portugués, Barros, Crujeiras, González-Manteiga, and Pereira (2014)) into our package. The addition of conditional copulae (Patton 2006; Fermanian and Wegkamp 2012) would also allow tackling issues, such as temporal autocorrelation, or performing time series analysis and regression.

## Supporting information

Code for replication

## Acknowledgements

JF received partial salary support from the Minnesota Agricultural Experimental Station. We thank Prof. Dr. Arpat Ozgul for his suggestions regarding automated copula fitting.

## A. Parametrizations of the wrapped Cauchy Distribution

The implementation of the wrapped Cauchy distribution in the **circular** package does not allow for the calculation of the distribution function or the quantile function. Therefore, we use a numerical approximation implemented in the **Wrapped** package. However, this package uses a different parametrization than then **circular** package. The equation for the wrapped Cauchy probability density with the parametrization of the **Wrapped** package is

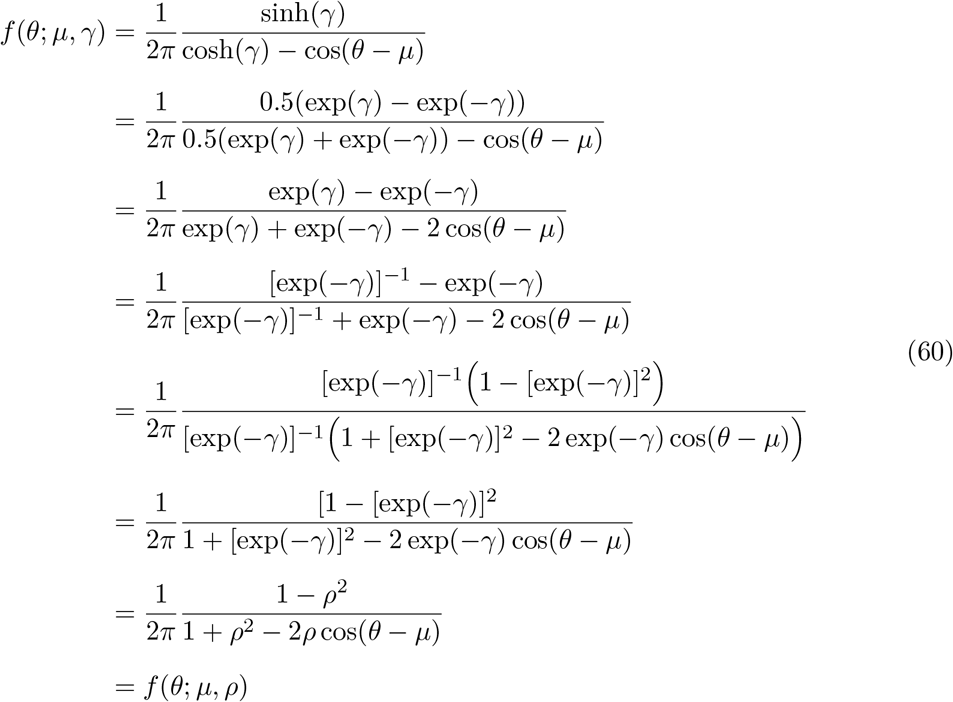

The last equality is in the parametrization of the wrapped Cauchy density of the **circular** package. The relationship between the two parametrizations is *ρ* = exp(–*γ*).

## B. Derivation of the von Mises copula CDF with Bessel functions

With the two substitutions *q* = 2*πs* and *r* = 2*πt*, we can write

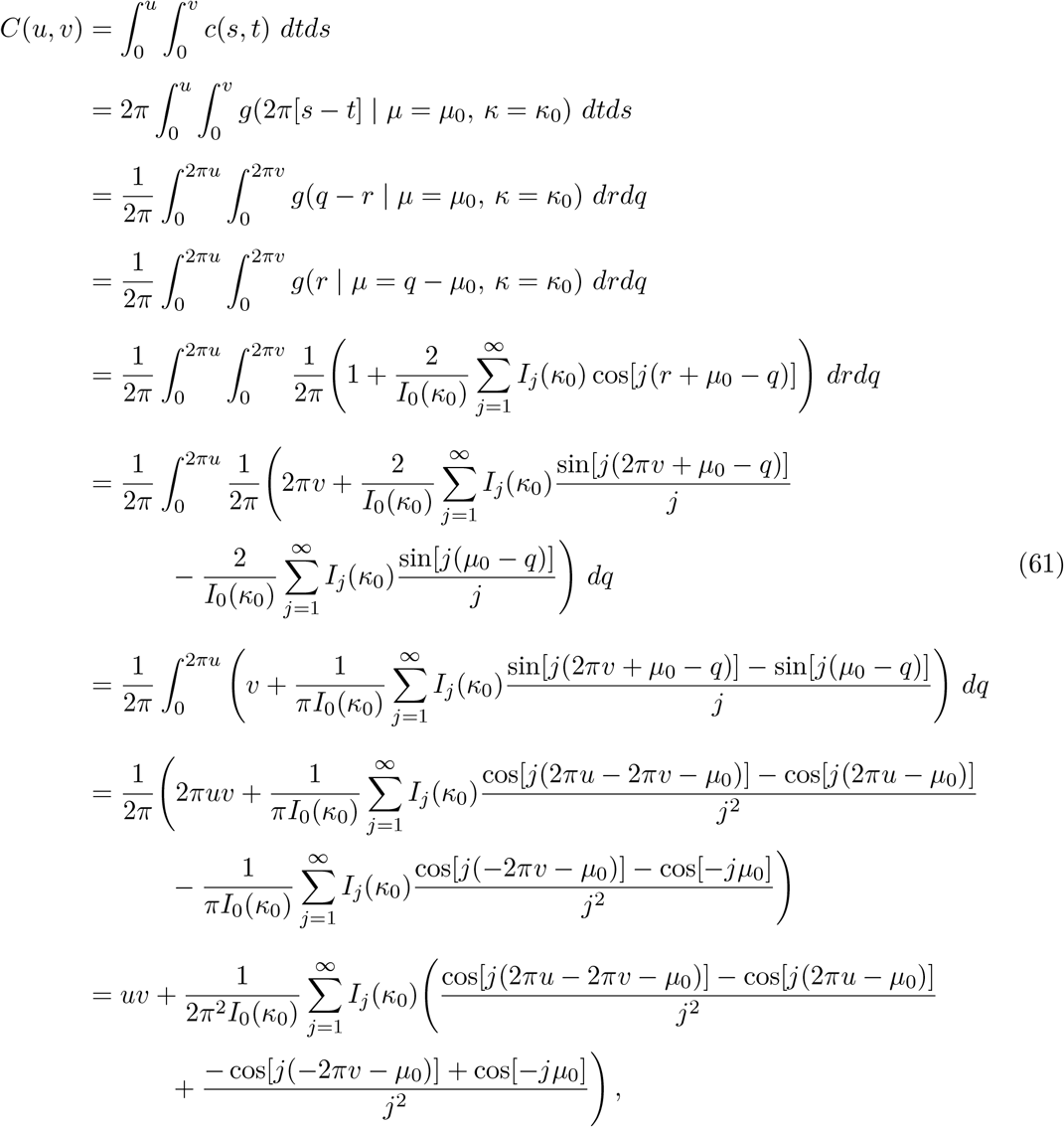

where *I_j_*(*x*) is the modified Bessel function of order *j*. Going from the third to the fourth line, we used the symmetry properties of *g*, which can be seen in Figure 2.

## C. Boundaries of copulae with quadratic sections

In Section 4.2.1, we stated the 4 necessary and sufficient conditions for a function *C*(*u*, *v*) = *uv* + *ψ*(*u*)*v*(1 – *v*) to be a copula with a density that is periodic in *u*. We chose *ψ*′(*u*) = *a*2*π* cos(2*πu*), which fulfills the second property, i.e. |*ψ*′(*u*)| ≤ 1 almost everywhere on [0, 1], only for –1/2*π* ≤ *a* ≤ 1/2*π*. The third property, i.e. |*ψ*(*u*)| ≤ min(*u*, 1 – *u*) ∀*u* ∈ [0, 1], can be rewritten as

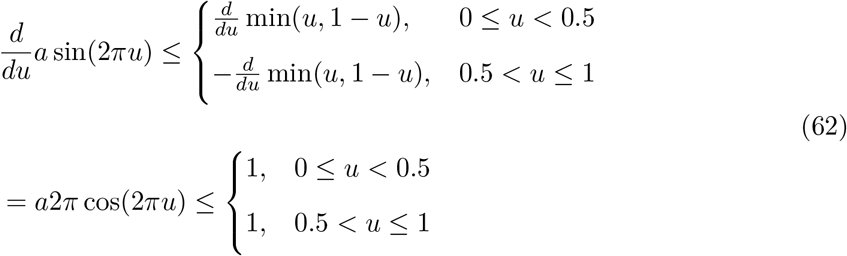

and is therefore also satisfied when – 1/2*π* ≤ *a* ≤ 1/2*π*.

## D. Boundaries of copulae with cubic sections

## E. Density of rectangular patchwork copulae

We can derive the density by taking the derivatives of the CDF with respect to *u* and *v*. First for (*u, v*) ∈ *R*_1_:

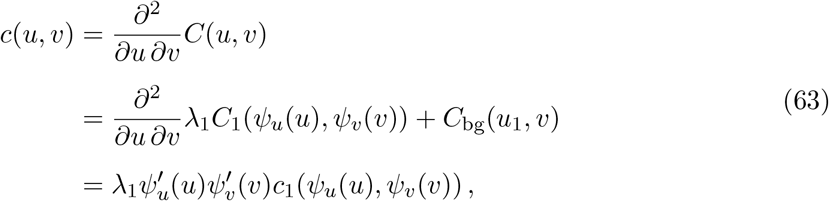

where we defined 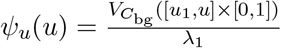 and 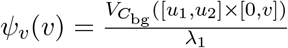. We now have to find the derivatives of these two functions

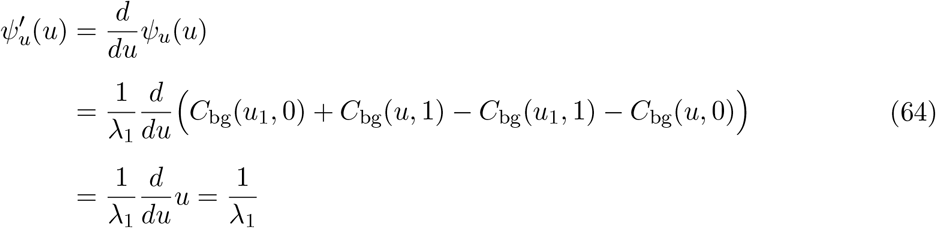

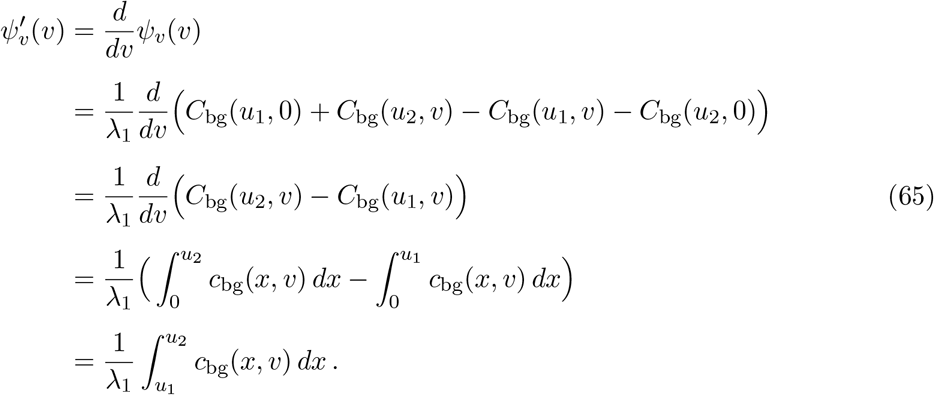

With similar calculations for *R*_2_ we finally arrive at Equation 54

## F. Perfect correlation

We obtained samples from a circular-linear, symmetric copula with perfect correlation using

~~~
*R> sample <- rcylcop(1000000, cyl_rect_combine(normalCopula(1)))*
~~~

These draws are shown in panel a of Figure 10. When we calculate the circular-linear correlation coefficient of this sample (which consists of points along the lines *u* = *v*/2 and *u* = 1 – *v*/2), we will obtain a value close to 1. The same is, however, not true for the normalized mutual information. Therefore, we have included the argument `symmetrize` in the function `mi_cyl()`. If `symmetrize = TRUE`, the empirical copula is calculated from the data and all *u*-values larger than 0.5 are set to 1 – 0.5. The result of `mi_cyl()` is then approximately 1 (for computational reasons it is actually only exactly 1 with very large sample sizes) in the case of perfect correlation (see panel b of Figure 10).

**Figure 10:**
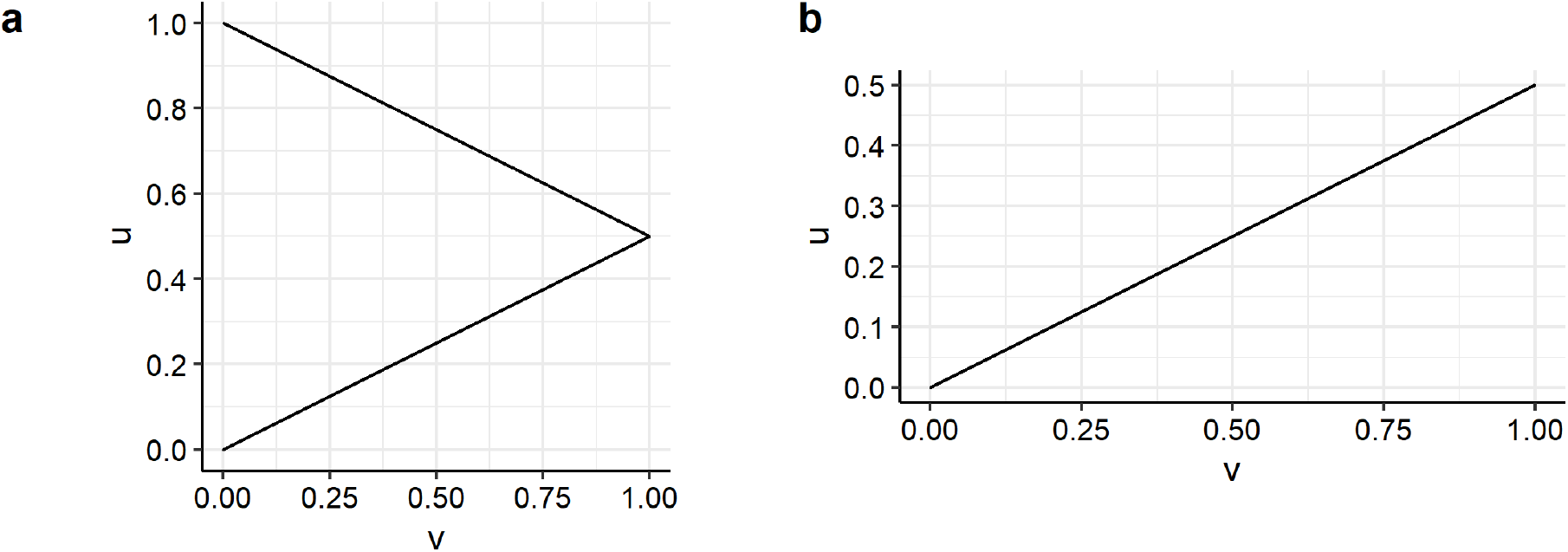
Left: draws from a circular-linear, symmetric copula with perfect correlation. Right: *u*-values larger than 0.5 of those draws are set to 1 – 0.5.

## G. Copulae considered in opt_auto()

The copulae that are fit to the data when using the `opt_auto()` function are

- `cyl_vonmises(flip = FALSE)`
- `cyl_vonmises(flip = TRUE)`
- `cyl_quadsec()`
- `cyl_cubsec()`
- `cyl_rot_combine(frankCopula(), shift = FALSE)`
- `cyl_rot_combine(claytonCopula(), shift = FALSE)`
- `cyl_rot_combine(gumbelCopula(), shift = FALSE)`
- `cyl_rot_combine(frankCopula(), shift = TRUE)`
- `cyl_rot_combine(claytonCopula(), shift = TRUE)`
- `cyl_rot_combine(gumbelCopula(), shift = TRUE)`
- `cyl_rect_combine(copula = frankCopula(), low_rect = c(0, 0.5), up_rect = “symmetric”, flip_up = TRUE)`
- `cyl_rect_combine(copula = claytonCopula(), low_rect = c(0, 0.5), up_rect = “symmetric”, flip_up = TRUE)`
- `cyl_rect_combine(copula = gumbelCopula(), low_rect = c(0, 0.5), up_rect = “symmetric”, flip_up = TRUE)`
- `cyl_rect_combine(copula = claytonCopula(), low_rect = c(0, 0.5), up_rect = “symmetric”, flip_up = FALSE)`
- `cyl_rect_combine(copula = gumbelCopula(), low_rect = c(0, 0.5), up_rect = “symmetric”, flip_up = FALSE)`

The rectangular patchwork copula based on the Frank copula with parameter *α* and `flip_up = FALSE` is the same as a rectangular patchwork copula based on the Frank copula with parameter –*α* and `flip_up = TRUE` and is therefore omitted from the list.

## H. Bandwidth and non-parameteric density estimates

We calculated non-parametric density estimates of angles drawn from a mixed von Mises distribution with parameters *μ*_1_ = 0. *μ*_2_ = *π, κ*_1_ = 2, *κ*_2_ = 1, and prop = 0.7. As bandwidth, we used 14.1 (`bw_1`) and 42.0 (`bw_2`), which we obtained with

~~~
*R> bw_1 <- opt_circ_bw(theta = traj$angle, loss=“adhoc”, kappa.est=“ML”)
R> bw_2 <- opt_circ_bw(theta = traj$angle,
+    loss=“adhoc”,
+    kappa.est=“trigmoments”
+  )*
~~~

. The resulting kernel density estimates are shown in Figure 11.

**Figure 11:**
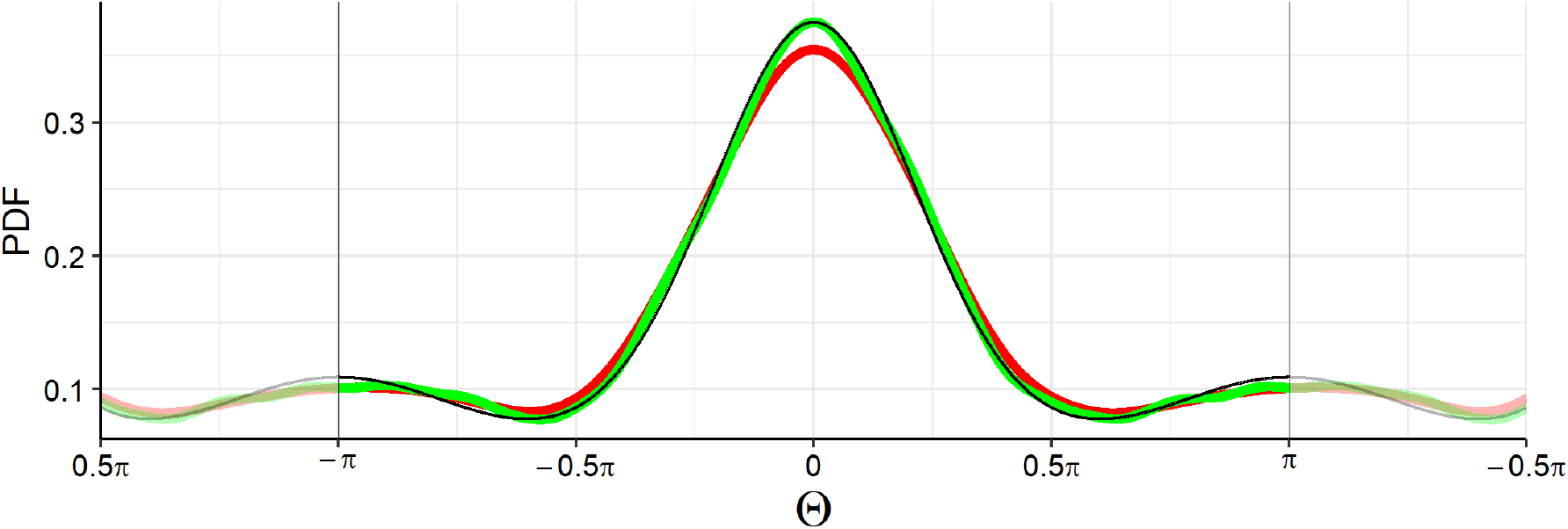
Black: true mixed von Mises density, red: non-parametric density estimate obtained with bandwidth 14.1, green: non-parametric density estimate obtained with bandwidth 42.0.

