## Supplementary material for "cylcop: An R Package for Circular-Linear Copulae with Angular Symmetry": Code for replication: replicate_paper.html

#### 2021-08-06

- Setup
- Figures in sections 1-6
  - Figure 1
  - Figure 2
  - Figure 3
  - Figure 4
  - Figure 5
- Coded examples
  - Figure 6
  - Figure 7
  - Results of section 6
  - Figure 8
- Appendix
  - Figure 10
  - Figure 11

### Setup

```
library(copula)
library(circular)
```

```
## 
## Attaching package: 'circular'
```

```
## The following objects are masked from 'package:stats':
## 
##     sd, var
```

```
library(cylcop)
```

```
## 
## Attaching package: 'cylcop'
```

```
## The following objects are masked from 'package:circular':
## 
##     dwrappedcauchy, rwrappedcauchy
```

```
library(cowplot)
library(ggplot2)
library(ggpubr)
```

```
## 
## Attaching package: 'ggpubr'
```

```
## The following object is masked from 'package:cowplot':
## 
##     get_legend
```

```
library(tidyverse)
```

```
## -- Attaching packages --------------------------------------- tidyverse 1.3.1 --
```

```
## v tibble  3.1.1     v dplyr   1.0.5
## v tidyr   1.1.3     v stringr 1.4.0
## v readr   1.4.0     v forcats 0.5.1
## v purrr   0.3.4
```

```
## -- Conflicts ------------------------------------------ tidyverse_conflicts() --
## x dplyr::filter() masks stats::filter()
## x dplyr::lag()    masks stats::lag()
```

```
cylcop_set_option(silent = TRUE)
```

### Figures in sections 1-6

#### Figure 1

```
outp <- data.frame(x = seq(-2 * pi, 2 * pi, 0.001))
outp$y <-
  map_dbl(outp$x,  ~ dvonmises(circular(.x), mu = circular(0.5 * pi), kappa =
                                 0.5))
x_breaks <- seq(-2 * pi, 2 * pi, 0.5 * pi)

plot_shift <- ggplot(outp[outp$x > -pi & outp$x < pi,], aes(x, y)) +
  geom_ribbon(
    aes(x = x, ymin = 0, ymax = y),
    alpha = 0.3,
    fill = "blue",
    linetype = 0
  ) +
  geom_line(size = 0.5) +
  scale_x_continuous(
    limits = c(-1.2 * pi, 1.2 * pi),
    breaks = x_breaks[3:7],
    labels = c(expression(-pi,-0.5 * pi, 0,
                          0.5 * pi, pi))
  ) +
  xlab(bquote(theta)) +
  ylab(bquote(f[Theta] ~ (theta))) +
  theme_bw() +
  theme(
    axis.title.x = element_text(size = 12, colour = "black"),
    axis.title.y = element_text(size = 12, colour = "black"),
    axis.text = element_text(size = 10, colour = "black"),
    panel.border = element_blank(),
    axis.line = element_line(colour = "black"),
    legend.position = "none",
    plot.margin = margin(10, 10, 10, 10)
  )

plot_zeroed <-
  ggplot(outp[outp$x > -0.5 * pi & outp$x < 1.5 * pi,], aes(x, y)) +
  geom_ribbon(
    aes(x = x, ymin = 0, ymax = y),
    alpha = 0.3,
    fill = "red",
    linetype = 0
  ) +
  geom_line(size = 0.5) +
  scale_x_continuous(
    limits = c(-0.7 * pi, 1.7 * pi),
    breaks = x_breaks[4:8],
    labels = c(expression(-0.5 * pi, 0,
                          0.5 * pi, pi, 1.5 * pi))
  ) +
  xlab(bquote(theta)) +
  ylab(bquote(f[Theta] ~ (theta))) +
  theme_bw() +
  theme(
    axis.title.x = element_text(size = 12, colour = "black"),
    axis.title.y = element_text(size = 12, colour = "black"),
    axis.text = element_text(size = 10, colour = "black"),
    panel.border = element_blank(),
    axis.line = element_line(colour = "black"),
    legend.position = "none",
    plot.margin = margin(10, 10, 10, 10)
  )


outp$y4 <-
  map_dbl(outp$x,
          ~ pvonmises(
            circular(.x),
            mu = circular(0.5 * pi),
            kappa = 0.5,
            from = circular(-0.5 * pi)
          ))
outp$y5 <-
  map_dbl(outp$x,
          ~ pvonmises(
            circular(.x),
            mu = circular(0.5 * pi),
            kappa = 0.5,
            from = circular(-pi)
          ))
outp[outp$x < -0.5 * pi,]$y4 <- 0
outp[outp$x > 1.5 * pi,]$y4 <- 1
outp[outp$x < -pi,]$y5 <- 0
outp[outp$x > pi,]$y5 <- 1
plot_cdf <- ggplot(outp, aes(x, y4)) +
  geom_line(size = 0.5, color = "red") +
  geom_line(data = outp, aes(x, y5), color = "blue") +
  scale_x_continuous(
    limits = c(-1.5 * pi, 2 * pi),
    breaks = x_breaks[3:8],
    labels = c(expression(-pi,-0.5 * pi, 0,
                          0.5 * pi, pi, 1.5 * pi))
  ) +
  xlab(bquote(theta)) +
  ylab(bquote(F[Theta] ~ (theta))) +
  theme_bw() +
  theme(
    axis.title.x = element_text(size = 12, colour = "black"),
    axis.title.y = element_text(size = 12, colour = "black"),
    axis.text = element_text(size = 10, colour = "black"),
    panel.border = element_blank(),
    axis.line = element_line(colour = "black"),
    legend.position = "none",
    plot.margin = margin(10, 10, 10, 10)
  )

fig_1 <-
  ggarrange(ggarrange(plot_shift, plot_zeroed, ncol = 2), plot_cdf, nrow = 2) %>%
  suppressWarnings()
fig_1
```

```
# ggsave(filename="fig_1.png",fig_1,dpi=300)
```

#### Figure 2

```
plot_periodic <- ggplot(outp, aes(x, y)) +
  geom_line(size = 0.5) +
  scale_x_continuous(
    limits = c(-2 * pi, 2 * pi),
    breaks = x_breaks,
    labels = c(
      expression(-2 * pi, -1.5 * pi, -pi,-0.5 * pi, 0,
                 0.5 * pi, pi, 1.5 * pi, 2 * pi)
    )
  ) +
  xlab(bquote(theta)) +
  ylab(bquote(g[Theta] ~ (theta))) +
  theme_bw() +
  theme(
    axis.title.x = element_text(size = 12, colour = "black"),
    axis.title.y = element_text(size = 12, colour = "black"),
    axis.text = element_text(size = 10, colour = "black"),
    panel.border = element_blank(),
    axis.line = element_line(colour = "black"),
    legend.position = "none",
    plot.margin = margin(10, 10, 10, 10)
  )
outp$y2 <-
  map_dbl(outp$x,
          ~ pvonmises(
            circular(.x),
            mu = circular(0.5 * pi),
            kappa = 0.5,
            from = circular(-pi)
          ))
outp$y3 <-
  map_dbl(outp$x,
          ~ pvonmises(
            circular(.x),
            mu = circular(0.5 * pi),
            kappa = 0.5,
            from = circular(-0.5 * pi)
          ))
plot_cdf_periodic <- ggplot(outp) +
  geom_line(aes(x, y2), size = 0.5, color = "blue") +
  geom_line(aes(x, y3), size = 0.5, color = "red") +
  scale_x_continuous(
    limits = c(-2 * pi, 2 * pi),
    breaks = x_breaks,
    labels = c(
      expression(-2 * pi,-1.5 * pi,-pi,-0.5 * pi, 0,
                 0.5 * pi, pi, 1.5 * pi, 2 * pi)
    )
  ) +
  xlab(bquote(theta)) +
  ylab(bquote(G[Theta] ~ (theta ~ '|' ~ l))) +
  theme_bw() +
  theme(
    axis.title.x = element_text(size = 12, colour = "black"),
    axis.title.y = element_text(size = 12, colour = "black"),
    axis.text = element_text(size = 10, colour = "black"),
    panel.border = element_blank(),
    axis.line = element_line(colour = "black"),
    legend.position = "none",
    plot.margin = margin(10, 10, 10, 10)
  )
fig_2 <-
  ggarrange(plot_periodic, plot_cdf_periodic, nrow = 2) %>% suppressWarnings()
fig_2
```

```
# ggsave(filename="fig_2.png",fig_2,dpi=300)
```

#### Figure 3

```
cop1 <- cyl_vonmises(mu = 0, kappa = 1)
cop2 <- cyl_vonmises(mu = pi, kappa = 4, flip = TRUE)
p1 <- cop_plot(cop1,
               type = "pdf",
               plot_type = "ggplot",
               resolution = 100)
```

```
p2 <- cop_plot(cop2,
               type = "pdf",
               plot_type = "ggplot",
               resolution = 100)
```

```
combine <-
  cowplot::plot_grid(p1,
                     p2,
                     ncol = 2,
                     labels = "auto",
                     label_size = 14) %>%
  suppressWarnings()
combine
```

```
# ggsave("fig_3.png", plot = combine,
#        width=7.43, height=2.74,
#        dpi = 300,device="png")
```

#### Figure 4

```
cop1 <- cyl_quadsec(a = 1 / (2 * pi))
cop2 <- cyl_cubsec(a = -1 / (2 * pi), b = 0.01)
p1 <- cop_plot(cop1,
               type = "pdf",
               plot_type = "ggplot",
               resolution = 100)
```

```
p2 <- cop_plot(cop2,
               type = "pdf",
               plot_type = "ggplot",
               resolution = 100)
```

```
combine <-
  cowplot::plot_grid(p1,
                     p2,
                     ncol = 2,
                     labels = "auto",
                     label_size = 14) %>%
  suppressWarnings()
combine
```

```
# ggsave("fig_4.png", plot = combine,
#        width=7.43, height=2.74,
#        dpi = 300,device="png")
```

#### Figure 5

```
cop1 <- cyl_rot_combine(rotCopula(claytonCopula(8), flip = TRUE))
cop2 <-
  cyl_rot_combine(rotCopula(claytonCopula(8), flip = TRUE), shift = TRUE)
cop3 <- cyl_rect_combine(rotCopula(claytonCopula(1.1), flip = TRUE))
cop4 <-
  cyl_rect_combine(
    rotCopula(claytonCopula(1.1), flip = TRUE),
    background = cyl_quadsec(),
    low_rect = c(0, 0.3)
  )
set.seed(123)
p1 <- cop_scat_plot(cop1)
p1
```

```
p2 <- cop_scat_plot(cop2)
p2
```

```
p3 <- cop_scat_plot(cop3)
p3
```

```
p4 <- cop_scat_plot(cop4)
p4
```

```
combine <- cowplot::plot_grid(
  p1,
  p2,
  p3,
  p4,
  ncol = 2,
  labels = "auto",
  label_size = 14,
  rel_widths = c(1, 1)
) %>%
  suppressWarnings()
# ggsave("fig_5.png", plot = combine,
#        width=7.43, height=2*2.74,
#        dpi = 300,device="png")
combine
```

### Coded examples

#### Figure 6

```
copula_1 <- cyl_cubsec(a = 0.05, b = -1 / (2 * pi))
copula_2 <-
  cyl_rect_combine(
    copula = copula::rotCopula(gumbelCopula(1.2), flip = TRUE),
    background = cyl_rot_combine(frankCopula(10), shift = FALSE),
    low_rect = c(0.1, 0.4),
    up_rect = c(0.6, 0.9),
    flip_up = TRUE
  )

p1 <-
  cop_plot(copula_1,
           type = "pdf",
           plot_type = "ggplot",
           resolution = 200)
```

```
p2 <-
  cop_plot(copula_2,
           type = "pdf",
           plot_type = "ggplot",
           resolution = 200)
```

```
## Warning in cop_plot(copula_2, type = "pdf", plot_type = "ggplot", resolution = 200): Maximum of pdf is 22.9; for better visulatization,
##  values larger than 3.44 (99.5 percentile) are cut off.
```

```
combine <-
  cowplot::plot_grid(p1,
                     p2,
                     ncol = 2,
                     labels = "auto",
                     label_size = 14) %>%
  suppressWarnings()
combine
```

```
# ggsave("fig_6.png", plot = combine,
#        width=7.43, height=2.74,
#        dpi = 300,device="png")
```

#### Figure 7

```
set.seed(123)

traj <- make_traj(
  n = 10000,
  copula = copula_1,
  marginal_circ = "mixedvonmises",
  parameter_circ = list(
    mu1 = 0,
    mu2 = pi,
    kappa1 = 2,
    kappa2 = 1,
    prop = 0.7
  ),
  marginal_lin = "gamma",
  parameter_lin = list(shape = 3)
)
p1 <- scat_plot(traj, periodic = TRUE)
p1
```

```
p2 <- traj_plot(traj)
```

```
p3 <- cop_scat_plot(traj)
p3
```

```
p4 <- circ_plot(traj)
```

```
combine <-
  cowplot::plot_grid(
    p1,
    p2,
    p3,
    p4,
    ncol = 2,
    labels = "auto",
    label_size = 14,
    rel_widths = c(1, 1)
  ) %>%
  suppressWarnings()
combine
```

```
 # ggsave("fig_7.png", plot = combine,
 #        width=7.43, height=2*2.74,
 #        dpi = 300,device="png")
```

#### Results of section 6

##### Correlation

```
cor_cyl(theta = traj$angle, x = traj$steplength)
```

```
## [1] 0.07490922
```

```
mi_cyl(
  theta = traj$angle,
  x = traj$steplength,
  normalize = TRUE,
  symmetrize = TRUE
)
```

```
## [1] 0.02147514
```

##### Automatic copula fitting

```
guess_fit <- opt_auto(theta = traj$angle, x = traj$steplength)
```

```
## Warning in opt_auto(theta = traj$angle, x = traj$steplength): For at least one
## copula, MLE did not converge!
```

```
guess_fit$copula[[1]]
```

```
## Cub. sect. copula 
## a = 0.05103531 
## b = -0.1591549
```

```
guess_fit$AIC[[1]]
```

```
## [1] -912.9146
```

```
guess_fit$copula[[2]]
```

```
## Quad. sect. copula 
## a = 0.1053691
```

```
guess_fit$AIC[[2]]
```

```
## [1] -758.4953
```

```
guess_fit$copula[[3]]
```

```
## Rectangular patchwork of Frank copula 
## alpha = 1.667804 
## low_rect1 = 0 
## low_rect2 = 0.5 
## up_rect1 = 0.5 
## up_rect2 = 1 
## upper copula is rotated 90 deg
## rectangles_symmetric: TRUE
```

```
guess_fit$AIC[[3]]
```

```
## [1] -747.0132
```

##### Manual copula fitting

```
start_val <- optCor(
  copula = cyl_cubsec(0, 0),
  theta = traj$angle,
  x = traj$steplength,
  acc = 0.01,
  n = 1000,
  method = "cor_cyl",
  parameter = "both"
)
start_val
```

```
## [1] -0.12915494  0.09084506
```

```
optimized <- optML(
  copula = cyl_cubsec(),
  theta = traj$angle,
  x = traj$steplength,
  parameters = c("a","b"),
  start = start_val,
  lower = c(-0.1, -1 / (2 * pi)),
  traceOpt = T,
  optim.method = "L-BFGS-B",
  optim.control = list(maxit = 100)
)
```

```
## param= -0.0999999985,  0.0908450569 => logL=  -1026.5782
## param= -0.0989999985,  0.0908450569 => logL=  -1019.6414
## param= -0.0999999985,  0.0908450569 => logL=  -1026.5782
## param= -0.0999999985,  0.0918450569 => logL=  -1035.6965
## param= -0.0999999985,  0.0898450569 => logL=  -1017.5237
## param=  0.159154941, -0.159154941 => logL=   195.60812
## param=  0.159154941, -0.159154941 => logL=   195.60812
## param=  0.158154941, -0.159154941 => logL=   206.23583
## param=  0.159154941, -0.158154941 => logL=   197.11853
## param=  0.159154941, -0.159154941 => logL=   195.60812
## param=  0.00741098224, -0.11513030857 => logL=   377.97467
## param=  0.00841098224, -0.11513030857 => logL=   379.46509
## param=  0.00641098224, -0.11513030857 => logL=   376.45827
## param=  0.00741098224, -0.11413030857 => logL=   376.14897
## param=  0.00741098224, -0.11613030857 => logL=   379.77632
## param=  0.0251209833, -0.1431492472 => logL=   438.04529
## param=  0.0261209833, -0.1431492472 => logL=   438.88196
## param=  0.0241209833, -0.1431492472 => logL=   437.18196
## param=  0.0251209833, -0.1421492472 => logL=   437.07941
## param=  0.0251209833, -0.1441492472 => logL=   438.98066
## param=  0.0466725173, -0.1591549407 => logL=   458.18119
## param=  0.0476725173, -0.1591549407 => logL=   458.29303
## param=  0.0456725173, -0.1591549407 => logL=   458.04068
## param=  0.0466725173, -0.1581549407 => logL=    457.9746
## param=  0.0466725173, -0.1591549407 => logL=   458.18119
## param=  0.0503851424, -0.1591549407 => logL=   458.45112
## param=  0.0513851424, -0.1591549407 => logL=   458.45552
## param=  0.0493851424, -0.1591549407 => logL=   458.41764
## param=  0.0503851424, -0.1581549407 => logL=   458.27919
## param=  0.0503851424, -0.1591549407 => logL=   458.45112
## param=  0.0510409395, -0.1591549407 => logL=    458.4573
## param=  0.0520409395, -0.1591549407 => logL=   458.44255
## param=  0.0500409395, -0.1591549407 => logL=   458.44288
## param=  0.0510409395, -0.1581549407 => logL=   458.29155
## param=  0.0510409395, -0.1591549407 => logL=    458.4573
## param=  0.0510352987, -0.1591549407 => logL=    458.4573
## param=  0.0520352987, -0.1591549407 => logL=   458.44271
## param=  0.0500352987, -0.1591549407 => logL=   458.44271
## param=  0.0510352987, -0.1581549407 => logL=   458.29149
## param=  0.0510352987, -0.1591549407 => logL=    458.4573
```

```
optimized
```

```
## $copula
## Cub. sect. copula 
## a = 0.0510353 
## b = -0.1591549 
## 
## $logL
## [1] 458.4573
## 
## $AIC
## [1] -912.9146
```

##### Parameteric and non-parametric estimation of the angular marginal distribution

```
#Estimation of the mixed von Mises parameters.
angle_distr <-
  fit_angle(theta = traj$angle,
            parametric = "mixedvonmises",
            mu = c(0, pi))
unlist(angle_distr$coef)
```

```
##       mu1       mu2    kappa1    kappa2      prop 
## 0.0000000 3.1415927 1.9633440 0.9550860 0.7109113
```

```
#Kernel density estimate of the circular marginal density.
bw <- opt_circ_bw(theta = traj$angle, loss = "adhoc", kappa.est = "trigmoments")
bw
```

```
## [1] 41.9904
```

```
angle_non_param <- fit_angle(theta = traj$angle, 
                               parametric = FALSE, 
                               bandwidth = bw
                             )
#Estimation of the gamma parameters.
step_distr <- fit_steplength(x = traj$steplength, parametric = "gamma")
unlist(step_distr$coef)
```

```
##     shape      rate 
## 2.9803921 0.9965143
```

```
#Produce a trajectory the estimated joint distribution described by the
#copula and the marginal distributions.
traj_fit <- make_traj(
  n = 10000,
  copula = optimized$copula,
  marginal_circ = "dens",
  parameter_circ = angle_non_param,
  marginal_lin = "gamma",
  parameter_lin = step_distr$coef
)
```

#### Figure 8

```
p1 <-
  cop_plot(
    guess_fit$copula[[2]],
    type = "pdf",
    plot_type = "ggplot",
    resolution = 200
  )
```

```
p2 <-
  cop_plot(
    guess_fit$copula[[3]],
    type = "pdf",
    plot_type = "ggplot",
    resolution = 200
  )
```

```
combine <-
  cowplot::plot_grid(
    p1,
    p2,
    ncol = 2,
    labels = "auto",
    label_size = 14,
    rel_widths = c(1, 1)
  ) %>%
  suppressWarnings()
combine
```

```
# ggsave("fig_code_3.png", plot = combine,
#        width=7.43, height=2.74,
#        dpi = 300,device="png")
```

### Appendix

#### Figure 10

```
set.seed(1)
sample <- rcylcop(100000, cyl_rect_combine(normalCopula(1)))
mi_cyl(
  theta = sample[, 1],
  x = sample[, 2],
  normalize = T,
  symmetrize = F
)
```

```
## [1] 0.80244
```

```
mi_cyl(
  theta = sample[, 1],
  x = sample[, 2],
  normalize = T,
  symmetrize = T
)
```

```
## [1] 0.9614457
```

```
sample <- as.data.frame(sample[1:10000,])
```

##### Plotting

```
plot_theme <- list(
  geom_point(size=0.01,alpha=0.5),
  theme(legend.position = "none"),
  theme_bw(),
  xlab("v"),
  ylab("u"),
  theme(
    axis.title = element_text(size = 12, colour = "black"),
    axis.text = element_text(size = 10, colour = "black"),
    panel.border = element_blank(),
    axis.line = element_line(colour = "black"),
    legend.position = "none"
  )
)
p1 <-
  ggplot(sample, aes(x = .data$v, y = .data$u)) +
  scale_y_continuous(breaks = seq(0, 1, 0.2)) +
  plot_theme +
  coord_fixed()

sample_sym <- sample
sample_sym$u <- modify_if(sample$u, ~ .x > 0.5, ~ 1 - .x)

p2 <-
  ggplot(sample_sym, aes(x = .data$v, y = .data$u)) +
  scale_y_continuous(breaks = seq(0, 1, 0.1)) +
  plot_theme +
  coord_fixed()

combine <- cowplot::plot_grid(p1,p2,ncol=2, labels="auto",label_size = 14,rel_widths = c(1,1))
combine
```

```
# ggsave("fig_appendix_2.png", plot = combine,
#        width=7.43, height=2.74,
#        dpi = 300,device="png")
```

#### Figure 11

```
bw_1 <- opt_circ_bw(theta = traj$angle, loss="adhoc", kappa.est="ML")
```

```
## Warning in opt_circ_bw(theta = traj$angle, loss = "adhoc", kappa.est = "ML"): ad-hoc method of finding bandwidth parameter might give very bad results, especially for multimodal population distributions
## you can use(kappa.est="trigmoments" or an mle method (loss="KullbackLeibler")
```

```
bw_1
```

```
## [1] 14.10933
```

```
bw_2 <- opt_circ_bw(theta = traj$angle, loss = "adhoc", kappa.est = "trigmoments")
bw_2
```

```
## [1] 41.9904
```

```
angle_non_param_1 <- fit_angle(theta = traj$angle, 
                               parametric = FALSE, 
                               bandwidth = bw_1
)
angle_non_param_2 <- fit_angle(theta = traj$angle,
                               parametric = FALSE,
                               bandwidth = bw_2
)

true_dens <- suppressWarnings(dmixedvonmises(as.numeric(angle_non_param_1$x),mu1 = 0,
                                             mu2 = pi,
                                             kappa1 = 2,
                                             kappa2 = 1,
                                             prop = 0.7))
```

##### Plotting

```
angle_non_param <-
  data.frame(x = c(
    as.numeric(angle_non_param_1$x),
    as.numeric(angle_non_param_2$x),
    as.numeric(angle_non_param_1$x)
  ),
  y = c(
    as.numeric(angle_non_param_1$y),
    as.numeric(angle_non_param_2$y),
    true_dens
  ))
angle_non_param$type <-
  c(rep(1, length(angle_non_param_1$x)), 
    rep(2, length(angle_non_param_2$x)),
    rep(3, length(angle_non_param_1$x)))
angle_non_param$type <- as.factor(angle_non_param$type)
angle_period_right <-
  data.frame(x = c(
    tail(
      as.numeric(angle_non_param_1$x),
      length(angle_non_param_1$x) / 4
    ) - (2 * pi),
    tail(
      as.numeric(angle_non_param_2$x),
      length(angle_non_param_2$x) / 4
    ) - (2 * pi),
    tail(
      as.numeric(angle_non_param_1$x),
      length(angle_non_param_1$x) / 4
    ) - (2 * pi)
  ),
  y = c(tail(
    as.numeric(angle_non_param_1$y),
    length(angle_non_param_1$y) / 4
  ),
  tail(
    as.numeric(angle_non_param_2$y),
    length(angle_non_param_2$y) / 4
  ),
  tail(
    true_dens,
    length(true_dens) / 4
  )))
angle_period_left <-
  data.frame(x = c(
    head(
      as.numeric(angle_non_param_1$x),
      length(angle_non_param_1$x) / 4
    ) + (2 * pi),
    head(
      as.numeric(angle_non_param_2$x),
      length(angle_non_param_2$x) / 4
    ) + (2 * pi),
    head(
      as.numeric(angle_non_param_1$x),
      length(angle_non_param_1$x) / 4
    ) + (2 * pi)
  ),
  y = c(head(
    as.numeric(angle_non_param_1$y),
    length(angle_non_param_1$y) / 4
  ),
  head(
    as.numeric(angle_non_param_2$y),
    length(angle_non_param_2$y) / 4
  ),
  head(
    true_dens,
    length(true_dens) / 4
  )))
angle_period_right$type <-
  as.factor(c(rep(1, length(angle_non_param_1$x) / 4), 
              rep(2, length(angle_non_param_2$x) / 4),
              rep(3, length(angle_non_param_1$x) / 4)
  ))
angle_period_left$type <-
  as.factor(c(rep(1, length(angle_non_param_1$x) / 4), 
              rep(2, length(angle_non_param_2$x) / 4),
              rep(3, length(angle_non_param_1$x) / 4)
  ))

p1 <- ggplot(angle_non_param,
             aes(
               x = as.numeric(.data$x),
               y = as.numeric(.data$y),
               group = .data$type
             )) +
  geom_line(aes(color = .data$type,size=.data$type)) +
  geom_line(
    data = angle_period_right,
    aes(
      x = as.numeric(.data$x),
      y = as.numeric(.data$y),
      group = .data$type,
      color = .data$type,
      size=.data$type
    ),
    alpha = 0.3
  ) +
  geom_line(
    data = angle_period_left,
    aes(
      x = as.numeric(.data$x),
      y = as.numeric(.data$y),
      group = .data$type,
      color = .data$type,
      size=.data$type
    ),
    alpha = 0.3
  ) +
  theme_bw() +
  xlab(bquote(Theta)) +
  ylab("PDF") +
  geom_vline(xintercept = c(-pi, pi),
             colour = "black",
             size = 0.2) +
  theme(
    axis.title.x = element_text(size = 17, colour = "black"),
    axis.title.y = element_text(size = 12, colour = "black"),
    axis.text = element_text(size = 10, colour = "black"),
    panel.border = element_blank(),
    axis.line = element_line(colour = "black"),
    legend.position = "none",
    plot.margin = margin(10, 20, 10, 10)
  ) +
  scale_x_continuous(
    breaks = seq(-1.5 * pi, 1.5 * pi, 0.5 * pi),
    limits = c(-1.5 * pi, 1.5 * pi),
    expand = c(0, 0),
    labels = c(expression(0.5 * pi,-pi,-0.5 * pi,
                          0,
                          0.5 * pi,
                          pi,-0.5 * pi))
  ) +
  scale_size_manual(values=c(1.5,1.5,0.5))+
  scale_color_manual(values = c("red", "green","black"))

p1
```

```
# ggsave("fig_appendix_3.png", plot = p1,
#        width=7.43, height=2.74,
#        dpi = 300,device="png")
sessionInfo()
```

```
## R version 4.0.5 (2021-03-31)
## Platform: x86_64-w64-mingw32/x64 (64-bit)
## Running under: Windows 10 x64 (build 19042)
## 
## Matrix products: default
## 
## locale:
## [1] LC_COLLATE=German_Switzerland.1252  LC_CTYPE=German_Switzerland.1252   
## [3] LC_MONETARY=German_Switzerland.1252 LC_NUMERIC=C                       
## [5] LC_TIME=German_Switzerland.1252    
## 
## attached base packages:
## [1] stats     graphics  grDevices utils     datasets  methods   base     
## 
## other attached packages:
##  [1] forcats_0.5.1   stringr_1.4.0   dplyr_1.0.5     purrr_0.3.4    
##  [5] readr_1.4.0     tidyr_1.1.3     tibble_3.1.1    tidyverse_1.3.1
##  [9] ggpubr_0.4.0    ggplot2_3.3.3   cowplot_1.1.1   cylcop_0.1.0   
## [13] circular_0.4-93 copula_1.0-1   
## 
## loaded via a namespace (and not attached):
##   [1] colorspace_2.0-0        ggsignif_0.6.1          ellipsis_0.3.1         
##   [4] rio_0.5.26              fs_1.5.0                rstudioapi_0.13        
##   [7] farver_2.1.0            gsl_2.1-6               fansi_0.4.2            
##  [10] mvtnorm_1.1-1           lubridate_1.7.10        xml2_1.3.2             
##  [13] knitr_1.33              jsonlite_1.7.2          broom_0.7.6            
##  [16] dbplyr_2.1.1            GoFKernel_2.1-1         stabledist_0.7-1       
##  [19] shiny_1.6.0             compiler_4.0.5          httr_1.4.2             
##  [22] backports_1.2.1         assertthat_0.2.1        Matrix_1.3-2           
##  [25] fastmap_1.1.0           lazyeval_0.2.2          cli_2.5.0              
##  [28] later_1.2.0             htmltools_0.5.1.1       tools_4.0.5            
##  [31] gtable_0.3.0            glue_1.4.2              movMF_0.2-5            
##  [34] Rcpp_1.0.6              carData_3.0-4           slam_0.1-48            
##  [37] cellranger_1.1.0        jquerylib_0.1.4         vctrs_0.3.7            
##  [40] crosstalk_1.1.1         xfun_0.22               rbibutils_2.1.1        
##  [43] openxlsx_4.2.3          rvest_1.0.0             mime_0.10              
##  [46] miniUI_0.1.1.1          lifecycle_1.0.0         rstatix_0.7.0          
##  [49] MASS_7.3-53.1           scales_1.1.1            hms_1.0.0              
##  [52] promises_1.2.0.1        yaml_2.2.1              curl_4.3               
##  [55] gridExtra_2.3           sass_0.3.1              stringi_1.5.3          
##  [58] highr_0.9               pcaPP_1.9-74            boot_1.3-27            
##  [61] zip_2.1.1               manipulateWidget_0.10.1 Rdpack_2.1.1           
##  [64] rlang_0.4.10            pkgconfig_2.0.3         rgl_0.106.8            
##  [67] evaluate_0.14           lattice_0.20-41         labeling_0.4.2         
##  [70] htmlwidgets_1.5.3       tidyselect_1.1.0        magrittr_2.0.1         
##  [73] R6_2.5.0                generics_0.1.0          ADGofTest_0.3          
##  [76] DBI_1.1.1               pillar_1.6.0            haven_2.4.1            
##  [79] foreign_0.8-81          withr_2.4.2             abind_1.4-5            
##  [82] pspline_1.0-18          modelr_0.1.8            crayon_1.4.1           
##  [85] car_3.0-10              KernSmooth_2.23-18      utf8_1.2.1             
##  [88] plotly_4.9.3            rmarkdown_2.7           viridis_0.6.0          
##  [91] grid_4.0.5              readxl_1.3.1            data.table_1.14.0      
##  [94] infotheo_1.2.0          reprex_2.0.0            digest_0.6.27          
##  [97] webshot_0.5.2           xtable_1.8-4            httpuv_1.6.0           
## [100] extraDistr_1.9.1        numDeriv_2016.8-1.1     stats4_4.0.5           
## [103] munsell_0.5.0           viridisLite_0.4.0       bslib_0.2.4
```
